# Systems biology analysis of vasodynamics in mouse cerebral arterioles during resting state and functional hyperemia

**DOI:** 10.1101/2025.05.06.652356

**Authors:** Hadi Esfandi, Mahshad Javidan, Eric R. McGregor, Rozalyn M. Anderson, Ramin Pashaie

## Abstract

Cerebral hemodynamics is tightly regulated by arteriolar vasodynamics. In this study, a systems biology approach was employed to investigate how the interplay between passive, myogenic, neurogenic, and astrocytic responses shapes arteriolar vasodynamics in small rodents. A model of neurovascular coupling is proposed in which neurons inhibit and dampen the myogenic response to promote vasodilation during activation, and facilitate the myogenic response to promote rapid vasoconstriction immediately post-activation. In this model, inhibition of the myogenic response is mediated by the hyperpolarization of smooth muscle and endothelial cells. Dampening and facilitation of the response are mediated by neuronal production of nitric oxide and release of neuropeptide Y, respectively. We also introduce a model for gliovascular coupling, in which astrocytes periodically inhibit the myogenic response upon detecting an increase in myogenic activity through interactions between their endfeet and arterioles. Our study revealed that in the resting state, the interplay between the delayed myogenic response and passive distension, acting as negative and positive feedbacks respectively, generates undamped oscillations in vessel diameter, known as vasomotion. In the active state, these oscillations are disrupted by the neurogenic and astrocytic responses. The biophysical model of arteriolar vasodynamics presented in this study lays the foundation for quantitative analysis of cerebral hemodynamics for cerebrovascular health diagnostics and hemodynamic neuroimaging.

**Author summary:** Cerebral hemodynamic imaging is widely used to investigate brain function in-vivo. These signals are primarily shaped by arteriolar vasodynamics, which result from a combination of physiological processes mediated by multiple interacting cell types. A biophysical model of this dynamics offers a valuable computational framework for achieving more accurate and quantitative interpretation of hemodynamic signals. In this study, I applied a computational biology approach to incorporate several well-established cellular signaling pathways into a unified model, which was used to investigate system-level arteriolar behavior and identify missing or less understood mechanisms involved in cerebral blood flow regulation. Our results show that arteriolar vasodynamics is not solely driven by neurogenic responses; astrocytic response and hemo-vascular interactions also play important roles in shaping the observed dynamics. The model also provided a means to explore how in-silico analysis of hemodynamic signals can reveal potential cellular-level impairments that manifest as system-level changes in cerebral hemodynamics. Incorporating our proposed biophysical model into cerebral hemodynamic analysis can improve the fidelity of hemodynamic imaging—enabling more accurate inference of regional neuronal and astrocytic activity from hemodynamic signals, and enhancing our ability to diagnose cerebrovascular pathologies.

## Introduction

Cerebral arteriolar vasodynamics regulates essential mechanisms of brain homeostasis, including oxygen and glucose delivery [1], neuronal function [2, 3], and glymphatic clearance [4]. While advances in in-vivo imaging techniques [5–8], genetic engineering [9], in-vitro vascular preparations [10, 11], and cellular manipulation techniques [12] have enriched the neurogliovascular coupling (NGVC) field, a biophysical model that elucidates the primary drivers of arteriolar vasodynamics has yet to be developed. In this article, we employed a computational and systems biology approach to study the dominant cell signaling pathways shaping arteriolar vasodynamics.

Functional hyperemia (FH) is a physiological process that boosts local blood flow in response to a local increase in brain activity. This increase in blood flow is facilitated by the transient vasodilation of feeding vessels mediated by the NGVC, and deformability of red blood cells (RBCs) within the brain’s vasculature [13]. We can categorize these interactions into two groups: feedforward response, and feedback response [14]. The feedforward response refers to cerebral blood flow (CBF) regulatory mechanisms where non-vascular cells, in proportion to their physiological demand, send signals to vascular cells to modulate the myogenic response and induce vasodilation. If the vasodilations induced by feedfoward responses fail to meet the demand during phases of intense and sustained activities, the feedback response, including the continuous feedback from metabolic deficiency sensors, is activated.

Research has demonstrated that RBCs act as oxygen sensors. RBCs autonomously regulate their deformability to modulate blood viscosity in capillaries in response to decreases in environmental oxygen tension [13]. We categorized the contribution of RBCs to FH as a feedback response, assuming their influence becomes significant only when vasodilation induced by feedforward NGVC cannot fully compensate for the oxygen consumed by activated brain cells. Moreover, biphasic vasodilations have been observed in the transitional zone (TZ) or pre-capillary arterioles during sustained FH, with the second delayed phase abolished by a K*_AT P_* channel blocker [15]. Based on these findings, we also categorized K*_AT P_* channel activation in vascular cells as a feedback response that is activated when the delivered blood cannot fully compensate for reduced nutrient levels. Recognizing the involvement of these feedback responses, along with other possible paths [16] [14], our study aimed to develop a model of the feedforward NGVC responses that shape arteriolar vasodynamics in a dynamically functioning mouse brain.

Arteriolar vasodynamics is largely driven by neuronal activity, with the reported correlation between the two estimated at approximately 60% in awake mice [17]. Also, astrocytes have been shown to contribute to CBF regulation [18, 19]. An open question is whether astrocytes dynamically modulate arteriolar vasodynamics and reduce this direct correlation. A recent study demonstrated that inhibiting astrocyte endfoot calcium signaling suppresses the delayed and secondary phases of sensory-evoked vasodilation in mouse penetrating arterioles (PAs) during sustained FH [12]. This finding suggests that astrocytes may regulate arteriolar vasodynamics conditionally, either by (1) inducing vasodilation in response to metabolic factors and/or (2) periodically initiating vasodilation at specific times. To investigate this conditional involvement, we employed a systems biology approach to disentangle neuronally from astrocyte-evoked vasodynamics and determine whether astrocyte-driven vasodynamics accounts for the portion of arteriolar vasodynamics uncorrelated to neuronal activity that observed in the awake mouse brain.

Vasodynamics is observed even in the absence of neuronal activity and is characterized by spontaneous low-frequency oscillations known as vasomotion. Vasomotion is considered a mechanism that facilitates efficient blood delivery and waste clearance [20–23]. Vasomotion is also observed in other organs, such as the skin [24], mesentery [25], and even in isolated arterioles [26]. The observation of vasomotion in isolated arterioles points to a potential vascular origin, although the underlying mechanisms are incompletely understood. A common strategy for generating sustained oscillations in biological regulatory systems involves the interplay between positive and delayed negative feedback loops [27, 28]. Most of the previously proposed vasomotion models have incorporated a cytosolic positive feedback into a Ca^2+^ release channel (be it either the ryanodine or the IP3 receptor) for sustaining oscillations [29, 30]. Here, we hypothesized that vasomotion may emerge from the interlinked positive and negative feedback loops, where passive distension amplifies delayed myogenic responses, leading to self-sustained oscillations in the cerebral vasculature. Using hemodynamic simulations in a simplified model of a network of coupled PAs, we investigate whether, in the absence of a cytosolic positive feedback loop, the model can still generate network-wide oscillations that resemble in-vivo vasomotion.

In our earlier work, we developed a coarsely segmented model of mouse cerebral vasculature, featuring a closed circulatory system and key morphological characteristics of mouse cerebrovasculature for in-silico analysis of static autoregulation [31]. In this study, we extend that framework to simulate hemodynamic and vasodynamic interactions within the vessel network under both active and resting brain states. This approach addresses a gap in existing computational NGVC studies by incorporating the often-overlooked influence of hemodynamics on vasodynamics. First, we build a cellular-level model of mouse PAs using a mathematical representation of their mechanobiological and electrophysiological properties and incorporate signaling pathways from other cell types that dynamically affect these properties (Section 1). We then use this PA model to simulate the kinetics of our proposed vasomotion (Section 2) and NGVC models (Section 3).

## Results

### Section 1: A Cellular-level Model of Mouse Penetrating Arteriole

In this study, we focused on PA vasodynamics for two reasons: 1) PAs are known to be the bottleneck of blood perfusion to the cortex [31–33], and 2) vasodynamic recordings of PAs are available in many studies, including those where NGVC was manipulated to silence a specific signaling pathway either genetically [9] or pharmacologically [12]. The myogenic response, regulated by hemodynamics and influenced by NGVC, is the primary driver of PA vasodynamics. Regulation of the myogenic response in PAs by hemodynamics is primarily adjusted by arteriolar SMCs (aSMCs) and, to a lesser extent, by arteriolar endothelial cells (aECs) [34]. In this section, we first design an endothelium-denuded segmented model of a mouse PA that incorporates an effective myogenic response by using a cellular-based model of aSMCs. Next, we add a simplified model of aECs to each PA segment and integrate this segmented vessel model into different PA segments within a larger cerebrovascular model.

The cerebrovascular model used in this study is an extension of our earlier work, where we designed an artificial cerebrovascular model in the form of a graph-based network composed of interconnected segmented cylindrical tubes with specific length and diameter [31] (Fig. 1), determining the resistance to blood flow according to Poiseuille’s law [35]. Briefly, we assumed that a typical PA in a mouse brain extends 840 *µ*m into the tissue, bifurcating at various depths to supply capillaries, with deeper segments accommodating less blood flow and having smaller diameters than superficial ones, as informed by Murray’s minimum-cost hypothesis [36]. Accordingly, the diameter of a PA, consisting of 28 segments, each 30 *µ*m long, was linearly reduced from 18 *µ*m at the surface to 12.6 *µ*m in the deepest cortical layers. In the following sections, we first model a vessel segment at 250 *µ*m depth with a maximum active diameter of 16.4 *µ*m and then extend this vessel segment model to all PA segments to create a PA model.

**Fig 1.**
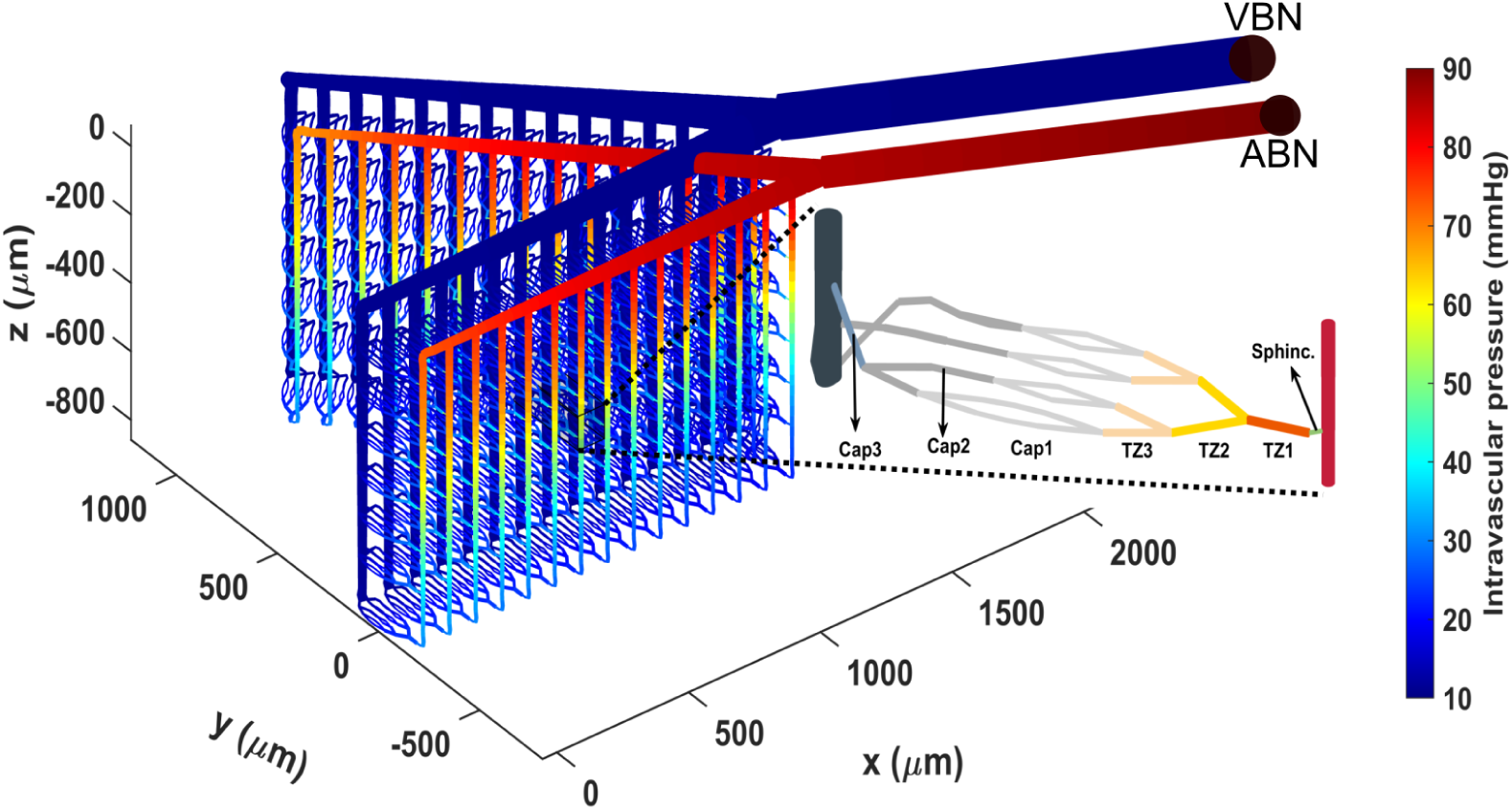
Designed segmented cerebral microvasculature model which includes two boundary nodes (ABN: artery boundary node, VBN: vein boundary node), a pial arteries network, and 30 PAs. The zoomed-in area highlights the structure of microvessels, including a sphincter, three layers of TZ segments (up to the 3rd order), and three layers of capillaries (beyond the 3rd order). The color coding illustrates the pressure distribution across the vascular segments, with boundary node pressures, ABNP and VBNP, set at 90 and 10 mmHg, respectively.

#### Endothelial-Denuded Segmented Model of Mouse Penetrating Arterioles

The diameter of a vessel segment is the result of the balance between two opposing processes: passive distension and active constriction. The magnitude of passive distension depends on the vessel’s distensibility and is modulated by circumferential wall tension (WT) [37]. The magnitude of active constriction in each vessel segment depends on the density of surrounding mural cells and is regulated by specific biomechanical forces exerted by the blood on the vessel wall. The exact biomechanical forces and mechanisms through which vascular cells (mural cells and endothelial cells) sense these forces and subsequently modulate their ionic channels are not fully understood, but are thought to mitigate the destructive forces of increased WT and wall shear stress (WSS).

In our earlier study, within the hypothetical autoregulation range for the designed cerebrovascular model, where the artery boundary node pressure (ABNP) varies between 40 and 130 mmHg while the vein boundary node pressure (VBNP) is held constant at 10 mmHg (Fig. 1), the constriction force of mural cells is linearly potentiated by the vessel WT [31, 38]. According to Laplace’s law, this tension is directly proportional to the IP within that segment and its internal diameter (*D*) and inversely proportional to the segment’s wall thickness (Δ), expressed as 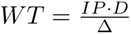. We did not account for the cytoskeletal remodeling, which likely occurs in pathological conditions like hypertension [37], and assumed that wall thickness remains constant under varying ABNP values. We quantified WT in all PA segments at ABNP = 130 mmHg (*WT*_max_) and used these values to model a linear relationship between SMC ion channel modulation and WT within the autoregulation range.

Myogenic tone (MT) development is primarily governed by SMC depolarization, activation of voltage-operated calcium channels (VOCCs), and an increase in intracellular calcium concentration [37]. Cell depolarization occurs either through the potentiation of depolarizing currents or the reduction of hyperpolarizing currents. Multiple ion channels have been implicated in the depolarization of SMCs (for review, see references [37] and [14]). Informed by the literature, our proposed SMC model includes WT-activated ion channels, such as TRPC6 [39], TRPM4 [40], and Ca^2+^-activated Cl*^−^* channels (TMEM16A) [41, 42], which play key roles in potentiating depolarizing currents in response to increase in WT, and WT-deactivated ion channels, including Kir channels [43, 44], which inhibit hyperpolarizing currents as WT increases.

Fig. 2 presents a schematic of the proposed aSMC-aEC model. Briefly, we assumed a linear relationship between the concentration of certain mechanotransduction signaling molecules (inositol trisphosphate (IP3) and diacylglycerol (DAG) [39]), and the exerted WT that potentiates depolarizing channels. This assumption is supported by several studies showing that increasing IP stimulates the phospholipase C (PLC) activity, which subsequently increases the concentration of second messengers (IP3 and DAG) in SMCs [45, 46], with subsequent upregulation of ion channels (TRPC6, TRPM4, and TMEM16A) to potentiate depolarizing currents. Fig. 3(a) depicts the relationship between the concentration of these second messengers and WT in our model, and for Kir channels, we simply assumed that their open probability (Kir_OP_) is directly downregulated by WT.

**Fig 2.**
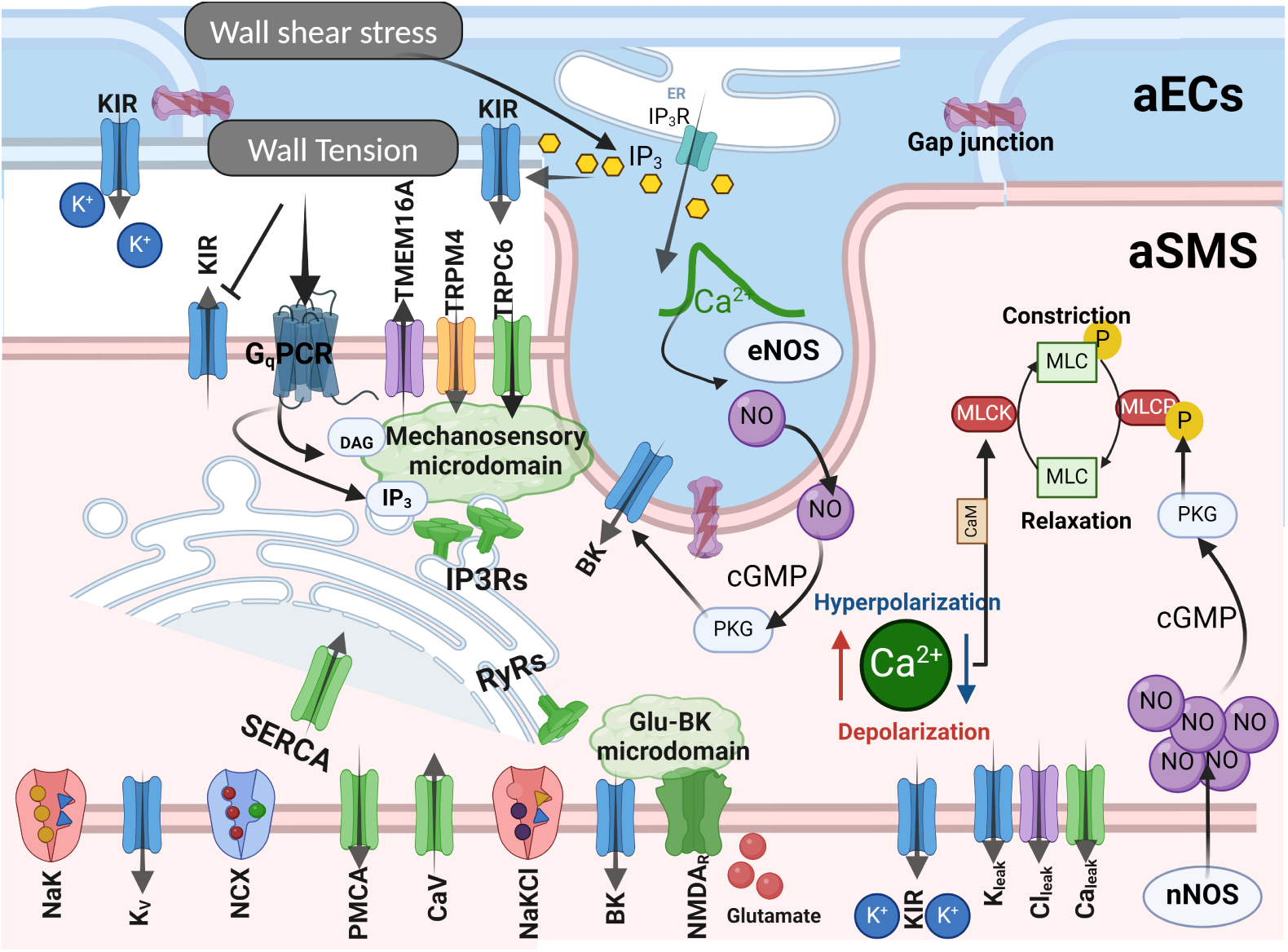
A schematic of the proposed SMC-EC model. For a detailed description we refer readers to the Methods section.The figure illustrates the channels and pumps regulating intracellular ion concentration and membrane potential in aSMCs, with color coding: potassium channels (blue), calcium channels and pumps (green), chloride channels (purple), and sodium channels (orange). The model includes two microdomains where calcium concentration can exceed the cytoplasm calcium concentration in aSMCs. The first is the mechanosensory microdomain, containing transient receptor potential (TRP) channels (e.g., TRPC6, TRPM4) and calcium-activated chloride channels (TMEM16A), which produce depolarizing currents in response to WT. In this microdomain, SMC mechanosensors are assumed to linearly convert WT into downstream signaling molecules, such as IP3 and DAG, modulating depolarizing currents. The second microdomain is the Glu-BK microdomain, where NMDA receptors, activated by neuron-released glutamate, trigger calcium efflux that activates nearby BK channels. The open probability of aSMC Kir channels is reduced by increasing WT, while larger WSS raises IP3 concentration within aECs, increasing both aEC Kir channel open probability and eNOS activity. The steady-state open probability of aSMC BK channels is increased by eNOS-dependent increases in NO/cGMP. In aSMC constriction mechanics, the rapid modulation of MLCK and MLCP activity is primarily driven by changes in cytoplasm SMC Ca^2+^ concentration and nNOS-dependent increases in NO/cGMP during increased neural activity, respectively.

**Fig 3.**
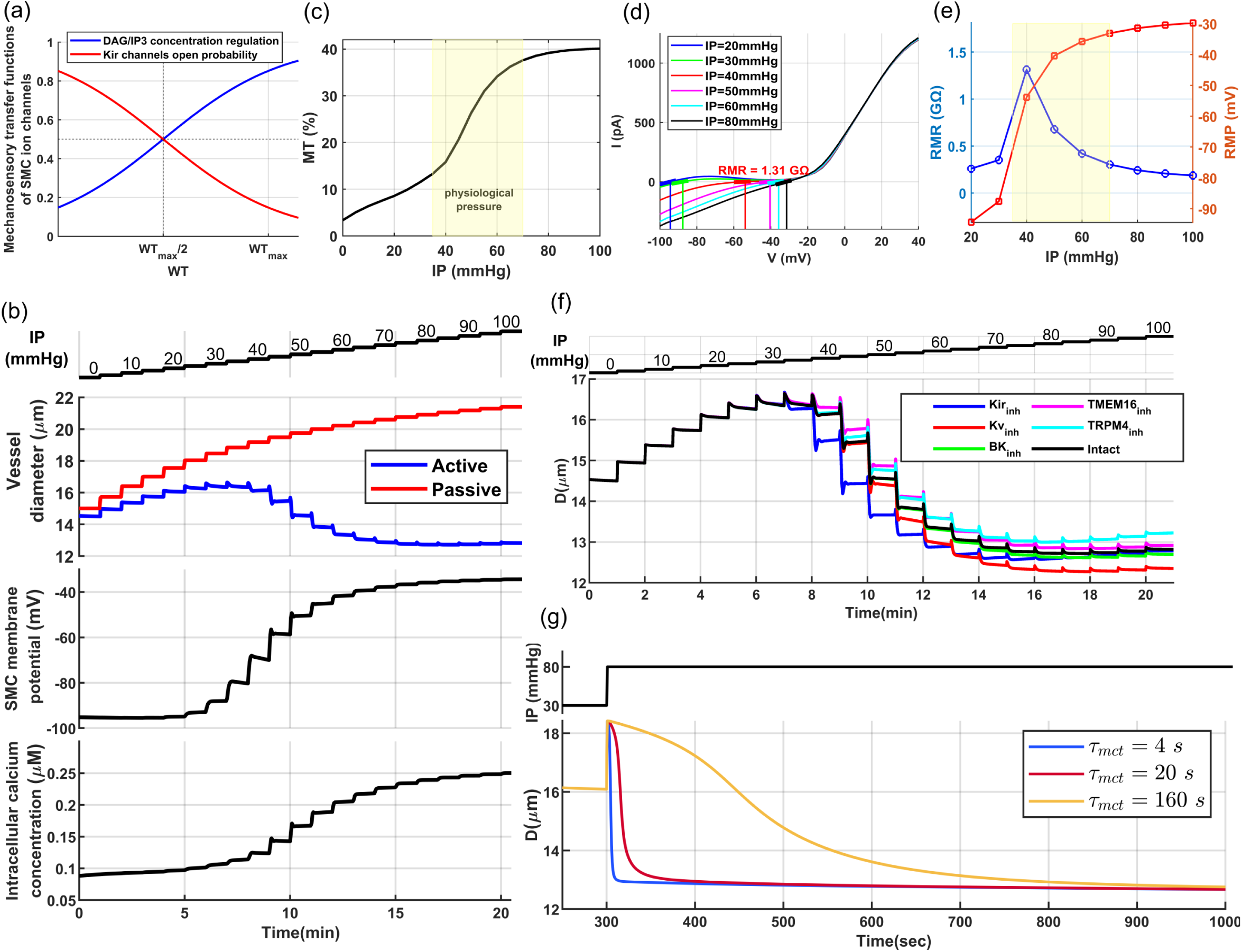
Endothelial-denuded segmented model of mouse PAs: (a) Quasi-linear sigmoidal function model the relationship between WT and mechanotransduction signaling molecules (IP3, DAG) and the WT-Kir_OP_ signaling pathway. (b) Pressure myography maneuver conducted on an endothelium-denuded vessel segment to calculate MT. (c) MT calculated for this segment, based on the curves plotted in panel (d-top), using 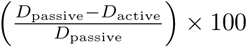. The physiological IP value within this segment ranges from 35–70 mmHg [34]. (d) The aSMC I-V curve under various IP values from simulated voltage-clamp analyses. Vertical lines depict the RMP where membrane current is near zero. The reciprocal of the slope of the tangent lines at the intersections of the vertical lines and curves indicates RMR. (e) Calculated RMR and RMP at different IP values. (f) Pressure myography test simulated under the assumption of 50%-inhibition of the main ion channels involved in MT regulation. (g) Simulated vasodynamics of the segmented model in response to a sudden increase in IP from 30 to 80 mmHg, analyzed under various values of the mechanotransduction time constant (*τ_mct_*).

We evaluated the performance of our proposed endothelial-denuded segmented PA model through a series of experiments before integrating it into the cerebrovascular model. Initially, we simulated a pressure myography test to evaluate MT development in our model through stepwise increases in IP within a PA segment under calcium-free (passive) and nominal external calcium concentration (active) conditions (Fig. 3(b)). Model parameters for passive distension were set so that the maximum passive diameter at high IP values reached about 130% of the maximum active diameter of the vessel segment [34]. For the active response, as expected, pressure elevation prompted membrane potential depolarization, leading to increased intracellular calcium concentration and subsequent vessel constriction (Fig. 3(b)).

Under low IP conditions (0-20 mmHg), WT-dependent depolarizing channels are less active, and a high open probability of Kir channels maintains the membrane potential near the equilibrium potential of potassium. Under these conditions, intracellular [Ca^2+^] is low, aSMCs are less capable of constricting the vessel, and the vessel passively distends until depolarizing channels become more active with increased WT, while simultaneously the open probability of Kir channels decreases. This leads to cell depolarization and a subsequent increase in [Ca^2+^] to a level that induces active constriction. The maximum active diameter of this vessel segment is achieved within the IP range of 30-35 mmHg. This IP range is in agreement with our cerebrovascular model design, where a PA segment at the depth of 250 *µ*m reaches its maximum dilation at the IP value between 30-35 mmHg when ABNP is set to 40 mmHg, which is the lower limit of the autoregulation range. Consistent with the in-silico and in-vivo analysis of autoregulation [31, 47], vessels must initially constrict steeply after reaching their maximum dilation to be at the optimal point of autoregulation, and as pressure increases, even a small constriction generates significant resistance to blood flow, optimizing the IP distribution in the network at the upper limit of the autoregulation range (Fig. 3(d)). The simulated values of membrane potential and [Ca^2+^] at various IP levels in our model (Fig. 3(b)) are closely aligned with values observed in in-vitro experiments for PAs [48–50], and the calculated MT of the modeled vessel segment (Fig. 3(c)) is consistent with the experimental data for PAs in both the maximum tone and profile [34, 51], indicating that the active constriction and passive distension models are effectively simulating the myogenic response of a PA vessel segment.

Next, we examined the resting membrane potential (RMP) and resting membrane resistance (RMR) which are critical electrophysiological parameters of mural cells. We simulated several voltage clamp tests under various pressure levels and we measured the steady-state membrane current while changing the SMC membrane potential from −100 to 40 mV (Fig. 3(f)). The generated I-V curves are in good agreement with the electrophysiology profile of freshly isolated SMCs from mouse brain arterioles [43, 44, 52]. At low IP values (unpressurized vessel in a physiological context), the high open probability of Kir channels and reduced dominance of WT-depolarizing channels result in an I-V curve for the SMC model that shows the typical characteristics of Kir channels [53, 54] including the negative slope and inward rectification. As IP increases, and within the physiological pressure range, the I-V curve of the cell has a shallow slope (reflecting low ion permeability) and maintains a high RMR, which is a characteristic of excitable cells. RMR reaches its maximum value at the lower limit of the physiological range, and as IP increases, RMR decreases and RMP rises (Fig. 3(d,e)), indicating that basal IP can modulate both RMP and RMR in SMCs, thereby affecting the sensitivity of the SMC membrane potential to external stimuli such as changes in hemodynamics or membrane potential of aECs.

In vascular SMCs, specific potassium channels operate under physiological conditions, mediating potassium efflux to counteract WT-dependent depolarizing currents, providing negative feedback for MT regulation [54], and maintaining a large RMR in aSMCs to enable rapid responsiveness to external stimuli. Arteriolar SMCs predominantly express four types of potassium channels [54]: ATP-sensitive (K*_AT P_*), large conductance Ca^2+^-activated (BK), inward rectifier (Kir), and voltage-gated (KV). It was suggested that K*_AT P_* channels play a minor role in cortical arterioles [16]. Furthermore, pressure-induced depolarization in arteriole SMCs triggers an amplification of Ca^2+^ sparks due to the facilitated Ca^2+^-induced Ca^2+^ release (CICR) process in ryanodine receptors (RyRs), driven by increased Ca^2+^ efflux via adjacent T-type VOCC [55], forming a negative feedback loop with coupled BK channels that mitigates myogenic vasoconstriction by promoting SMC hyperpolarization. However, unlike cortical surface arteries, BK channel-mediated K^+^ efflux does not inhibit MT significantly in PAs, and under physiological conditions, only Kir and KV channels are highly activated in aSMCs [54]. Consequently, in our proposed SMC model, TMEM16A and TRPM4 channels are key depolarizing channels, while Kir, KV, and BK channels serve as key hyperpolarizing channels, with a less contribution from the latter.

To evaluate the contribution of these channels to SMC constriction, we simulated a pressure myography test under the assumption of 50% channel inhibition (Fig. 3(f)). Kir channel inhibition significantly disrupted MT regulation within the physiological IP range. These simulation results align well with in vitro findings, where vessels exposed to a Kir channel blocker under physiological IP were significantly constricted [43], suggesting that SMC Kir channels play a crucial role in cerebral autoregulation. KV and BK channel inhibition caused MT dysregulation at mid to high IP values due to their voltage-dependent activation mechanism, with BK channels playing a much lesser role in the negative feedback regulation of MT in endothelium-denuded PAs. Inhibition of both key depolarizing channels, TRPM4 and TMEM16A, led to reduced vessel constriction.

Although the steady-state SMC mechanotransduction signaling pathways (such as DAG, IP3, and Kir_OP_) are regulated by WT, the cerebral vasculature operates in a dynamic environment where WT continuously fluctuates across vessel segments due to events like FH or changes in mean arteriolar pressure. Because mechanotransduction in SMCs does not occur instantaneously, there is a time delay in the adjustment of WT-dependent ion channel activity in response to rapid changes in WT. We modeled this delay using an exponential function with a mechanotransduction time constant (*τ*_mct_). This time constant plays a crucial role in shaping the temporal dynamics of the myogenic response, influencing the kinetics of FH and vasomotion, as explored in the following sections.

Various studies affirm that Neuropeptide Y (NPY) can enhance the vasoconstrictor actions of other molecules [56–59]. This facilitating effect is attributed to the synergistic interactions between NPY receptors (NPYRs) and other G-protein-coupled receptors (GPCRs) involved in vasoconstriction [60]. This synergy could potentiate transduction pathways following the mechanoactivation of Gq-coupled receptors in SMCs and lower the *τ*_mct_. We estimate that under physiological conditions, the value of *τ*_mct_ could range between 1 to 20 seconds and will provide evidence to support this. However, in-vitro conditions may result in much larger *τ*_mct_ values and a myogenic response that is several orders of magnitude slower than under physiological conditions [34, 37, 51, 61]. Facilitation of the myogenic response by NPY is particularly relevant in cerebral arterioles, as NPY receptors are highly expressed in aSMCs, and there are direct NPY interneuron–aSMC junctions [62]. Fig. 3(g) shows vessel diameter changes in response to an abrupt increase in IP from 30 to 80 mmHg in our model under three different *τ*_mct_, where the vessel initially passively distends, and the time to reach steady state can vary significantly depending on the value of *τ*_mct_.

#### Fast-Acting Downregulation of SMC constriction in Endothelial-Denuded Vessel Segment

To simulate arteriolar vasodynamics, we need to identify and model the fast-acting downregulators of SMC constriction that induce rapid vasodilation during periods of increased neural activity. The regulation of SMC constriction involves two competing processes: myosin light chain (MLC) phosphorylation by myosin light-chain kinase (MLCK), counteracted by dephosphorylation through myosin light-chain phosphatase (MLCP). A decrease in intracellular calcium concentration inhibits calcium-calmodulin-dependent MLCK activation, thereby promoting muscle relaxation [63]. Additionally, MLCP is activated by protein kinase G (PKG) and MLCK is inhibited by protein kinase A (PKA), with PKG and PKA themselves being activated by secondary messengers cyclic guanosine monophosphate (cGMP) and cyclic adenosine monophosphate (cAMP), respectively. These secondary messengers are primarily produced by NO-dependent sGC [64] and hormone/neurotransmitter-stimulated adenylyl cyclase (AC) [65, 66], respectively.

SMC intracellular calcium concentration is predominantly regulated by VOCCs, and reducing the SMC membrane potential or directly inhibiting VOCCs are the primary mechanisms of lowering SMC [Ca^2+^] and MT inhibition. Based on the reported in-vivo SMC [Ca^2+^] [67], we ruled out the possibility that fast VOCC inhibition via NGVC affects SMC constriction state during FH. We only focused on modeling the mediators that can rapidly modulate SMC membrane potential. Modulation of the SMC membrane potential is a multi-faceted process which is affected by hemodynamic changes, synaptic-like transmission between neural axons and SMCs [62], myoendothelial gap junctions [10], NO pathways [68], acidosis [16], and direct extracellular interactions between ion channels, such as astrocyte endfeet BK channels and SMC Kir channels [69]. We ruled out the possibility of the brain tissue acidification during FH, but other pathways were modeled in this study.

Kir channels are key candidates for modulating SMC membrane potential via NGVC, as they play a crucial role in SMC membrane potential autoregulation (Fig. 3(f)).The vasodilatory response to the potassium released from astrocyte endfeet BK channels is mediated by SMC Kir channels [69–71]. Elevation of extracellular potassium increases Kir channel conductance which hyperpolarizes the cell. Fig. 4(a) shows the vasodilatory response of the PA segmented model to the elevation of extracellular potassium (*K*_ex_) at discrete levels for three IP and two SMC *τ*_mct_ values. At IP = 50 mmHg, extracellular potassium elevation induces the largest vasodilation compared to higher IPs. In this physiological IP range, the cell’s RMR is large (Fig. 3(d,e)), and slight variations in Kir channel conductance can change the SMC membrane potential significantly. At higher IP values, RMR and Kir channel open probability decrease, making Kir channels less capable of inducing hyperpolarization in the SMC.

**Fig 4.**
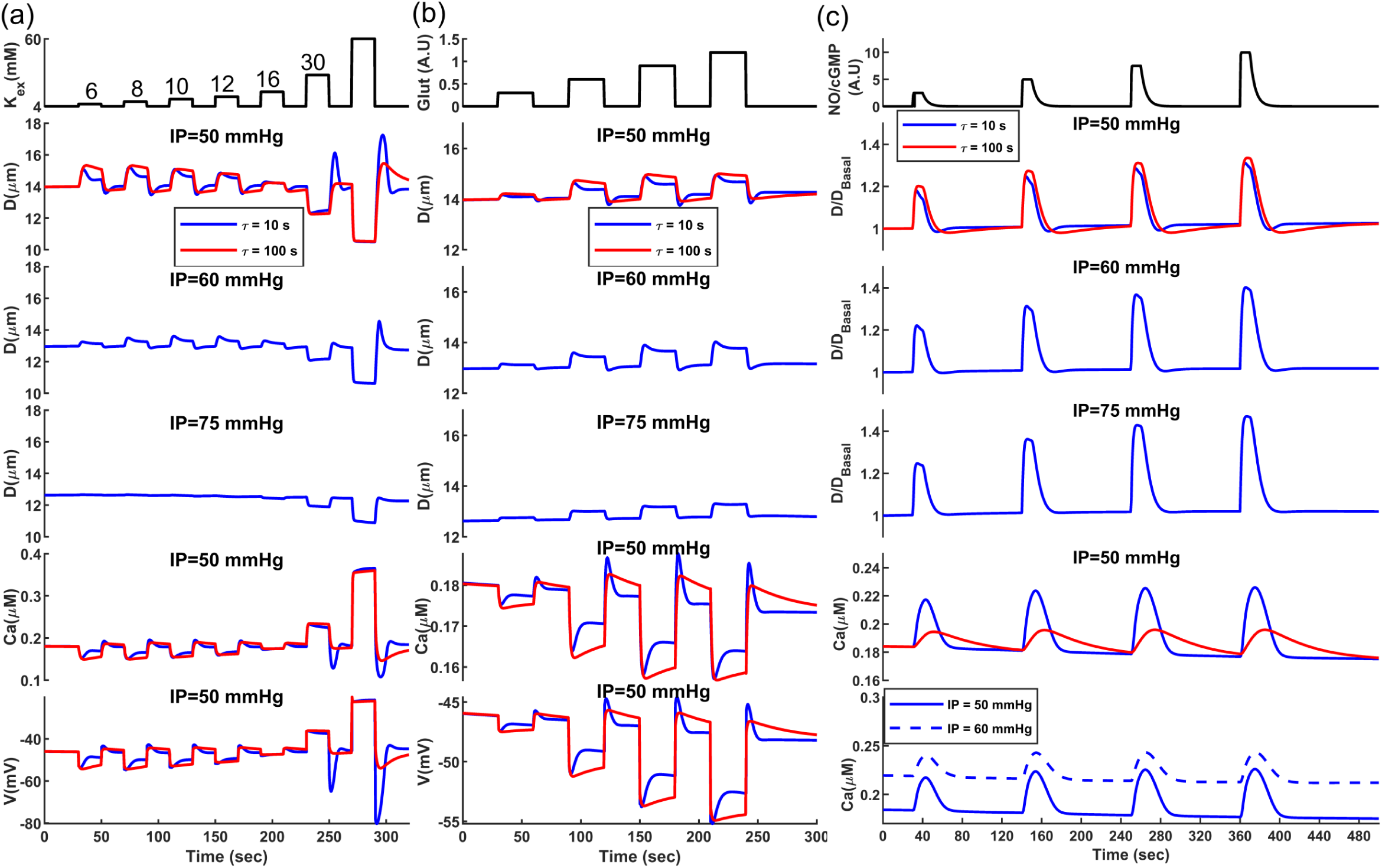
Analysis of vasodynamics in an endothelium-denuded model under the influence of vasodilatory mediators. Vasodilatory responses were measured for all mediators under two *τ*_mct_ and three IP values. (a) Changes in vasodynamics, SMC membrane potential, and [Ca^2+^] when increasing the SMC extracellular potassium concentration from a baseline of 4 mM up to 60 mM in discrete steps. (b) Changes in vasodynamics, SMC membrane potential, and [Ca^2+^] when increasing the glutamate concentration from a baseline of 0 to 1.2 AU in four discrete steps. (c) Changes in vasodynamics and [Ca^2+^] when increasing SMC NO/cGMP concentrations from a baseline of 0 to 10 AU in four discrete steps. Numbers considered for the concentration of glutamate and NO were hypothetical since the exact physiological values are unknown. Model parameters were adjusted accordingly to produce the maximum dilatory response at the highest hypothetical concentrations.

The SMC mechanotransduction time constant (*τ*_mct_) can also affect the magnitude of vasodilation. When *τ*_mct_ is small, SMCs respond rapidly to WT changes. The initial vasodilation increases WT according to Laplace’s law, leading to cell depolarization that counteracts Kir channel-induced hyperpolarization. This myogenic response is more pronounced at lower IP values because, under these conditions, the cell’s RMR is large, and slight changes in WT-dependent currents have a large impact on the SMC membrane potential.

At IP = 50 mmHg, the largest vasodilation was observed at [*K*^+^]_ex_ = 8 mM, but the peak shifts to higher potassium concentrations at IP = 60 mmHg because altering extracellular potassium not only impacts Kir channel conductance but also changes the potassium Nernst potential. Increasing IP also raises intracellular potassium concentration, further affecting the potassium Nernst potential [72]. Elevating extracellular potassium up to 16 mM induces vasodilation at IP = 50 and 60 mmHg, but it results in slight constriction at IP = 75 mmHg. Therefore, the direction and extent of diameter change depend on both extracellular potassium concentration and the value of IP [69]. At a high [*K*^+^]_ex_, the potassium Nernst potential is much larger than its physiological level, leading to a significant SMC depolarization (∼-20 mV at *K*_ex_ 60 mM) and causing maximum constriction in the vessel, which typically does not occur at any IP value under physiological extracellular potassium concentration.

BK channels are modulated by the membrane potential and Ca^2+^ sparks near the cell membrane, which makes them key candidates for fast hyperpolarization and relaxation of aSMCs [73]. Studies have shown that BK channel activation can induce pronounced vasodilation in PAs when exogenous activators are present. For example, brain tissue acidification dilated PAs rapidly due to a dramatic increase in Ca^2+^ spark activity and BK channel function, driven by the enhanced RyR open probability [16]. BK channels in PA SMCs are also colocalized with GluN1, a subunit of NMDA receptors, and glutamatergic neurons have been shown to dilate arterioles through synaptic-like transmission at neural–aSMC junctions (NsMJs). Activation of NMDA receptors in Glu-NsMJs leads to a Ca^2+^ ion influx, which binds to and activates BK channels, resulting in increased potassium ion efflux, membrane hyperpolarization, and subsequent relaxation of aSMCs. In-vivo disruption of Glu-NsMJs inhibits NGVC substantially [62]. To incorporate this NGVC mechanism, we included a Glu-BK microdomain in our model, where activation of glutamate receptors locally increases [Ca^2+^] in the microdomain and activates BK channels (Fig. 2).

Fig. 4(b) shows the vasodilatory response of the PA segmented model to elevated glutamate at four discrete hypothetical concentrations, across three IP and two SMC *τ*_mct_ values. The highest hypothetical glutamate concentration induces a 3.5% steady-state dilation at IP = 50 mmHg (after balancing neurogenic and myogenic responses when *τ*_mct_= 10 s), while this level is about 5% at IP = 60 mmHg. This occurs because BK channels are mediated by both Ca^2+^ and membrane potential. At IP = 60 mmHg, the higher SMC membrane potential enhances the vasodilatory effect of glutamate. The myogenic response following initial vasodilation is less pronounced at IP = 75 mmHg due to small RMR, where changes in WT-dependent currents are less capable of altering the membrane potential and the [Ca^2+^]. The post-stimulus undershoot in the diameter change (observed in the blue curves of Fig. 4(a,b)) has the same origin as the initial overshoot and is associated with the potentiation/inhibition of myogenic response following vasodilation/constriction. Once glutamate or [*K*^+^]_ex_ return to the baseline, the membrane potential is expected to return to its baseline. However, since MT is larger than its basal level in the dilation phase, following the vasoconstriction due to the removal of the stimulus, the delay in the adjustment of MT causes the vessel to further constrict, which produces a post-stimulus undershoot.

Nitric oxide (NO) is a potent vasodilator produced rapidly by both endothelial (eNOS) and neural nitric oxide synthase (nNOS) and plays a crucial role in NGVC [74]. In-vivo evidence shows that neuronally produced NO acts directly on arterioles [9, 75, 76]. Blockade of neural NO synthase has the most significant inhibitory effect on the feedforward mechanisms [74], which can be compensated by feedback mechanisms [77]. NO relaxes vascular SMCs and dilates blood vessels by increasing the intracellular cGMP concentration, which stimulates cGMP-dependent PKG activity. However, the rapid vasodilatory signalling pathway mediated by PKG is not fully understood. This complexity arises from the diverse ways that PKG can influence the SMC constriction state. PKG modulates various SMC ion channels, including the activation of BK channels [78], inhibition of TRPM4 channels [68], and inhibition of VOCCs [79], collectively leading to decreased SMC [Ca^2+^]. PKG also directly activates MLCP [80, 81]; however, it has remained unclear whether PKG-dependent MLCP activation or PKG-SMC ion channel modulation plays a more significant role during the rapid dynamics of FH. Here, we assume that the fast-acting effect of NO on aSMCs is primarily governed by the increased MLCP activity rather than by SMC ion channel modulation. This assumption is supported by the studies which show that the early phase of NO-induced vasodilation involves the suppression of CPI-17 phosphorylation which is an MLCP inhibitory protein [82–84]. Therefore, in our vessel wall mechanical model adopted from [85–87], variations in NO/cGMP/PKG activity directly affect the kinetics of actin-myosin interactions, ultimately leading to SMC relaxation.

Fig. 4(c) shows the vasodilatory response of the PA segmented model to elevated NO/cGMP at four discrete hypothetical levels, across three IP and two SMC *τ*_mct_ values. Despite the immediate rise in NO/cGMP concentration, we assumed a degradation rate for these molecules that delays their decline under physiological conditions. This degradation was modeled by a decaying exponential function with a 5-sec time constant, regardless of any dose dependency that might exist in reality. The relative diameter change of NO-induced dilation is larger at a higher basal IP. This occurs because the increase in MLCP activity correlates with a decrease in MLC phosphorylation sensitivity to [Ca^2+^] [82], resulting in [Ca^2+^] having a less pronounced effect on SMC contraction during periods of elevated NO/cGMP. Therefore, during NO/cGMP dose-dependent relaxation of SMCs, vessels with higher basal IP experience larger passive distention, leading to a more pronounced diameter change.

If the myogenic response is defined as the intrinsic ability of SMCs to modulate their membrane potential and [Ca^2+^] by WT, then increased MLCP activity due to the NO/cGMP/PKG pathway can be seen as a mechanism of dampening this response by reducing the sensitivity of MLC phosphorylation to MLCK activity. On the other hand, hyperpolarizing SMCs with mediators can be viewed as the inhibition of myogenic response. During NO-induced dilation, the increase in diameter and WT leads to cell depolarization, causing a rise in [Ca^2+^] and potentiation of the myogenic response (Fig. 4(c)). However, since the NO/cGMP/PKG pathway dampens the myogenic response, the effect of the potentiated myogenic response is small and negligibly counteracts the vasodilatory effect of NO. Comparing the NO vasodilatory response under two analyzed mechanotransduction time constants at IP = 50 mmHg shows that a smaller *τ*_mct_ leads to a sooner and more pronounced increase in [Ca^2+^], but this potentiation of the myogenic response during the dilatory phase only negligibly reduces the peak amplitude of NO-induced dilation compared to the case of a larger *τ*_mct_.

Rapid myogenic response potentiation (*τ*_mct_ =10 s) can help the vessel return to its basal level sooner after the stimulus; however, the slow degradation rate of NO/cGMP and the dampened myogenic response in the presence of these molecules does not lead to a pronounced post-stimulus undershoot. As NO/cGMP degrades and returns to its basal state, the myogenic response may cause a post-stimulus undershoot if [Ca^2+^] is above pre-stimulus levels. In the *τ*_mct_ =100 s case, where the myogenic response occurs more slowly, once NO/cGMP returns to its basal state and Ca^2+^ sensitivity of MLC phosphorylation increases, the post-stimulus undershoot period is longer. However, due to a smaller increase in myogenic response-induced [Ca^2+^], the degree of post-stimulus undershoot is only marginally larger than in the fast myogenic response scenario, where a significant rise in [Ca^2+^] occurs alongside the reduced Ca^2+^ sensitivity of MLC phosphorylation. Similar to the vasodilatory response of other mediators, like [*K*^+^]_ex_ and glutamate, the degree of post-stimulus undershoot decreases with increasing IP. This occurs because, under higher IP, cell RMR is small, and WT-dependent depolarizing currents have a reduced impact on the membrane potential, resulting in a smaller increase in [Ca^2+^] (Fig. 4 (c - bottom panel)). Consequently, smaller post-stimulus undershoots are observed at IP = 75 mmHg and IP = 60 mmHg compared to IP = 50 mmHg. In the following sections, as we analyze the neurogenic impulse response of the CBF regulatory system, we will present strong experimental evidence to support our analysis of the dynamics of NO-induced vasodilation.

#### Integrating Segmented Model of Mouse Penetrating Arterioles into Cerebral Vasculature Model

Thus far, we have developed a segmented model of an endothelium-denuded vessel and simulated the primary vasodilatory signaling pathways. Endothelial cells are also well recognized for their crucial role in FH dynamics. In this section, we propose a simplified model of aECs to be added to the PA segment model, which we then integrate into the PA segments of the cerebral vasculature to capture their physiological contribution to FH dynamics. ECs are known to serve as sensors of neural activity within the CBF regulatory system, relaying this information retrogradely up the vascular tree via electrical coupling [10]. Active neurons discharge potassium ions, which enhance the conductance of Kir channels in ECs, leading to fast hyperpolarization of arteriolar ECs in both proximal and distal regions of the activated area, which is transmitted to SMCs through MGJs and results in their dilation [88, 89].

ECs sense changes in potassium concentration and translate that to changes in their membrane potential. A large RMR, in the range of several Gigaohms [90, 91], makes ECs sensitive to such changes, allowing them to respond quickly and precisely. [53] proposed an abstract mathematical model for ECs, which includes a model for the Kir current (*I_Kir_*) while combining all other transmembrane currents into one nonspecific linear background current, *I_bg_*(*v*) = *G_bg_*(*V_EC_* − *E_bg_*), where *E_bg_* = −30 mV was assumed to be constant. In our proposed model, we replaced *I_bg_* with 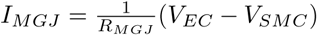. Here, *R_MGJ_* represents the resistance of gap junctions between an EC and SMC which was assumed to be 1 GΩ [53, 92]. This replaced depolarizing current opposes *I_Kir_*-induced hyperpolarization to maintain a large RMR in ECs, and couple the EC membrane potential with the membrane potential of mural cells for autoregulation.

Fig. 5(a) shows that, unlike the endothelium-denuded model, increasing the value of IP does not fully depolarize the SMC due to large Kir currents in coupled aEC. Consequently, SMC [Ca^2+^] does not exceed 150 nM, allowing passive distention to counteract the low contractile force of SMCs, which results in dilation at high IP values. In this pressure myography test, IP was increased with no blood flow in the vessel, resulting in zero WSS. The Kir current in aECs is also dependent on WSS, with WSS activating this channel [43]. At low mean arteriolar pressure, where blood flow is minimum and vessel diameter is maximum, WSS is minimum, nullifying the hyperpolarizing effect of EC Kir channels. As mean arteriolar pressure increases, WT-dependent currents in SMCs can constrict the vessel. As SMCs depolarize, the EC membrane potential also depolarizes due to gap junction coupling and small EC Kir channel activity under small WSS. With increasing SMC constriction force and vessel contraction, WSS rises which activates EC Kir channels; however, the EC is now less polarized due to coupled depolarized SMC. Therefore, Kir channel conductance is reduced at physiological extracellular potassium concentrations. Thus, EC Kir channel activity has a negligible effect on SMC membrane potential, allowing SMCs to autoregulate while maintaining EC Kir channels functional [43]. Increasing extracellular potassium of ECs by neuronally discharged potassium during FH causes a rightward and upward shift of the *I_Kir_* curve [53], inducing EC hyperpolarization, which in turn hyperpolarizes SMCs.

**Fig 5.**
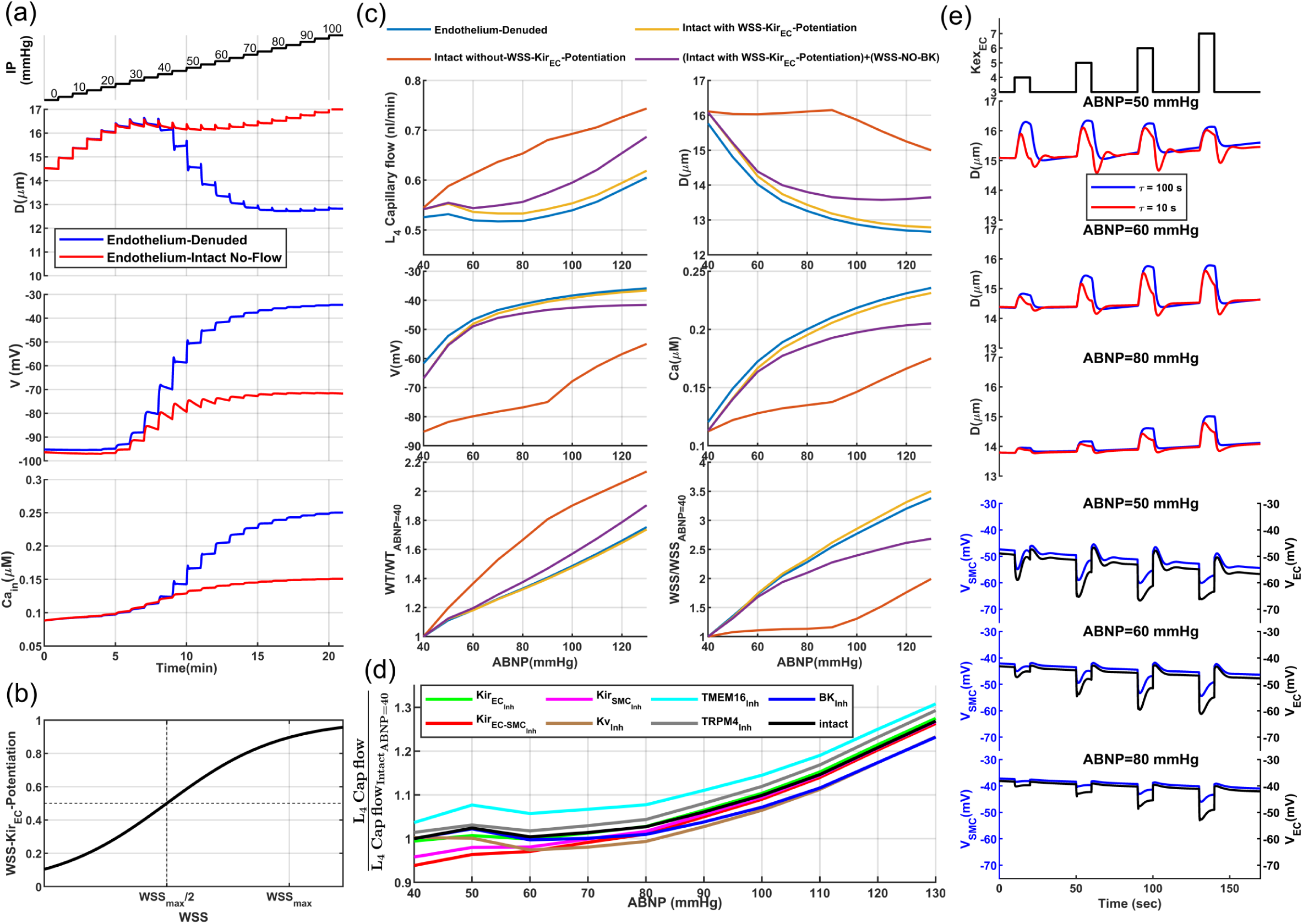
Integration of Intact segmented model of mouse PAs into the cerebral vasculature model: (a) pressure myography test performed on an intact vessel segment with no blood flow and exerted WSS. Note: these simulation results are not physiologically valid, as no WSS dependency for EC-Kir channel activation was considered in the model. In the absence of flow, EC-Kir channels are deactivated (panels b, c), and MT is likely correctly developed in vessels without flow [93]. (b) A quasi-linear sigmoidal function models the relationship between WSS and Kir*_EC_* open probability. (c) Analysis of the autoregulatory performance of the intact PA model. Purple curves are associated with a segment of the final PA model, where aEC respond to increased WSS caused by the rising ABNP. This response results in an increase in the open probability of EC Kir channels and the potentiation of eNOS-based signaling. NO produced by eNOS diffuses into the SMC, potentiating BK channels to enhance MT negative feedback and mitigate the elevated WSS. (d) Analysis of dysregulation with a 50% inhibition of primary ion channels in the PA model. (e) Vasodynamic analysis of the PA model under two different SMC *τ_mct_* and three ABNP values, in response to an increase in EC extracellular potassium concentration from a baseline of 3 mM up to 7 mM in four discrete steps.

To run simulations with this model, blood needs to flow into the vessel segment to exert WSS. This requires a circulatory network where blood flows through the network based on the IP gradient between its entry and outlet nodes. Taking a stepwise approach, we first integrated the endothelium-denuded segmented model of PA into the designed cerebral vascular model. Our earlier study demonstrated that, morphological differences exist between PA segments [31]. Superficial segments generally have larger diameters to accommodate higher blood flow and have higher expression of contractile elements to counteract passive distension for cerebral autoregulation. The relative expression of contractile elements in each segment was modeled by introducing a relative contractility index. Model parameters associated with the baseline diameter, passive distension, and expression of contractile elements (relative contractility) were tuned in the proposed endothelium-denuded segmented model, extending the segmented model to all 28 segments of each PA in the designed cerebral vasculature.

Once the endothelium-denuded PA model established, we set the VBNP to 10 mmHg and increased the ABNP from 40 to 130 mmHg in 10 mmHg increments, where PA segment diameters were automatically adjusted by the integrated endothelium-denuded model, while diameters of other segments, including surface arteries, sphincters, TZ, and capillaries, were manually optimized based on the myogenic response for each ABNP [31]. At ABNP = 130 mmHg, PA segments experience maximum WSS, as the blood flow peaks and vessel diameters are minimum. We quantified WSS values in all PA segments at ABNP = 130 mmHg and saved them as the maximum WSS (*WSS*_max_) of each segment. These *WSS*_max_ values were then used to model the modulation of EC Kir channel open probability in response to an imposed WSS in intact PA model, using a quasi-linear sigmoidal function(Fig. 5(b)). In this intact PA model, the homocellular gap junctions between adjacent aSMCs and aECs have a resistance of 100 megaΩ [90, 94, 95].

To evaluate the developed intact PA model, we analyzed its performance in autoregulation. In our previous work, we showed that with optimal MT potentiation in all segments of our cerebral vasculature model, when ABNP increases within the autoregulation range, blood flow in non-bifurcating capillaries (Cap1 in Fig. 1(a)) remains around 0.5 nl/min. We also demonstrated that MT in TZ vessels and sphincters primarily autoregulates superficial blood flow, while MT in PAs plays a more pronounced role in deeper layers [31]. Building on this, we calculated the steady-state values of deep capillary (*L*4_Cap1_) blood flow and other model variables for a PA segment at a depth of 250 *µ*m to assess the intact PA model’s autoregulation performance.

Adding the WSS-dependency of Kir channel activity to the endothelium-intact PA model (yellow curves) provides a similar autoregulatory performance to the endothelium-denuded model (blue curves) and also corrects the dysregulation observed in the absence of WSS-dependency in the EC Kir current (orange curves)(Fig. 5(c)). Within the physiological ABNP range of 40-80 mmHg (corresponding to 30-65 mmHg IP within PA), in scenarios where MT is correctly adjusted in vessels (blue and yellow curves), blood flow in L4 non-bifurcating capillaries remains relatively constant. When ABNP increases above this range, even though the myogenic induced constriction in PA segments could not sustain plateaued autoregulation, *L*4_Cap1_ blood flow moderately increases, which is in agreement with the computational and experimental observations [31, 47]. SMC membrane potential and [Ca^2+^] are also in the range reported in in-vitro studies [48]. The maximum MT level at ABNP = 130 mmHg is about 42% 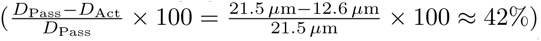 for the analyzed PA segment, which matches the reported maximum MT in PAs [34].

WSS-dependent activation of eNOS is also known to inhibit MT in vessels [96]. In mouse PAs, this NO-induced dilation has been linked to increased SMC BK channel activity [34]. We adjusted the model parameters associated with the WSS/NO/BK pathway to reduce *MT*_max_ by approximately 14% compared to the case without the WSS-eNOS pathway (decreasing from 42% to 36%). As expected, when the WSS-NO pathway is included (purple curves) and the vessel is less constricted at the upper end of the autoregulation range, *L*4_Cap1_ blood flow increases at a higher rate. Additionally, while WT increases slightly compared to the case of no WSS-NO-BK pathway (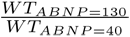 changes from 1.75 to 1.9), WSS increases much less in the presence of the WSS-NO pathway (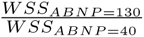 changes from 3.5 to 2.65). Within the autoregulation range, as mean arteriolar pressure increases, MT potentiation mitigates the WT increase associated with the rising ABNP, while MT inhibition via WSS mitigates the increased WSS. This regulation implies that vessels potentially adapt through both WT-dependent mitigation of WT via SMCs, and WSS-dependent mitigation of WSS via ECs, preventing excessive mechanical stress that could compromise vascular integrity and optimal function.

Next, we aimed to qualitatively demonstrate how disruption of different ion channels in our PA model can lead to cerebral dysregulation (Fig. 5(d)). The simulation tested a 50% inhibition of various ion channels in the PA model. Focusing on the PA, and optimally adjusting other segemnts’ MT for autoregulation, the most dysregulated scenario occurs when Kir channels are disrupted in vascular cells, as SMC Kir current inhibition leads to more constricted vessels within the physiological pressure range (Fig. 3(f)). Oxidative stress and inflammation in pathological conditions such as Alzheimer’s disease (AD) can impair Kir channel function [97–99], which might explain the hypoperfusion previously reported in AD mouse models [100, 101]. As expected, the inhibition of TMEM16A channels and KV channels results in the hyperperfusion and hypoperfusion, pointing to their crucial roles in MT adjustment at different WT levels in aSMCs. Inhibition of BK channels also led to less WSS-dependent inhibition of MT in the upper range, and consequently, a smaller increase in blood flow.

After evaluating the intact PA model, we aimed to demonstrate that its EC Kir channels are functional in converting neural activity into vessel dilation. We addressed the question whether active neurons discharge potassium around ECs in specific vascular zones to hyperpolarize the endothelial network through gap junctions, or if all parenchymal ECs experience increased extracellular potassium during neural activity. In both scenarios, it seems logical that the EC basal membrane potential is mediated by MT in mural cells to maintain the autoregulation process. An increase in aEC [*K*^+^]_ex_, leads to endothelial network hyperpolarization in proportion to the amount of potassium released, which is subsequently transferred to SMCs, inducing vasodilation (Fig. 5(e)). This proportionality is less evident in the case of ABNP = 50 mmHg due to the instability of both SMC and EC membrane potentials during the test, which continuously decline and impair the correlation between [*K*^+^]_ex_ and vasodilation. Under very low mean arterial brain pressure (ABNP = 50 mmHg), aSMCs operate at low IP (30-40 mmHg depending on cortical depth), small WT-depolarizing currents, and large RMR (Fig. 3(d, e)), which makes them sensitive to minor changes in the dominant SMC Kir currents. The negative slope region of the Kir channel I-V curve explains this continuous downward trend (Fig. 3(d) - green curve). A slight initial hyperpolarization can trigger a positive feedback loop, increasing Kir channel conductance and driving the membrane potential further toward negative values until the Kir current saturates at the peak of the negative slope region. This persistent hyperpolarization and vasodilation might serve as a protective mechanism under low mean arterial pressure, maximizes vessel dilation, and increases arterial pressure to counteract the low perfusion.

### Section 2: Modeling the Kinetics of Vasomotion

Vasomotion, characterized by low-frequency, undamped oscillations of arteriolar diameter, is a feature of cerebral vasculature dynamics observed during the resting state [102–104]. In this state, neural activity is limited to baseline levels, and we assumed that the myogenic response in arterioles is facilitated by the tonic release of mediators from various brain cells, including NPY and other signaling molecules.

A delayed negative feedback, including the delay in the myogenic response, can induce oscillations. When WT in a vessel segment increases, the myogenic response triggers the muscular constriction to counteract the elevated WT. Delayed readjustment of the muscular force generates an overshoot in the system’s response which deviates WT in the opposite direction. The repeating cycles of overshoots and delayed corrections generate oscillations (Fig. 6(a)). Variations in the vessel diameter cause changes in the exerted WT and hemodynamics, and with some delay, modulate the SMC membrane potential. Range-normalized curves reveal a 2 sec delay between the WT trough and the SMC membrane potential trough in our model (t = 18.5 sec vs. t = 20.5 sec (Fig. 6(a-right panel))). The adjustment of the SMC [Ca^2+^] via VOCCs lags behind changes in its membrane potential due to the time required for Ca^2+^ ions to enter or exit the cell until [Ca^2+^] is tuned to the membrane potential. In our model, this delay is about 1.5 sec (t = 20.5 sec vs. t = 22 sec). The delay in the adjustment of MLC phosphorylation in response to changes in [Ca^2+^] further delays the adjustment of SMC constriction and vessel diameter, resulting in a 1.7 sec delay between the vessel diameter and SMC [Ca^2+^] curves (t = 22 sec vs. t = 23.7 sec). This timing aligns with the time-lag observed in-vivo between vessel diameter and SMC [Ca^2+^] changes in awake mice [67]. Overall, there is a cumulative 5.2 sec delay between the moment the vessel segment experiences the minimum WT and the time the SMC constriction force reaches to its minimum which is the end of the dilation phase. A similar dynamic occurs in the opposite direction, but with faster Ca^2+^ dynamics during constriction, resulting in a 4.2 sec lag between the maximum WT and maximum SMC constriction force which is the end of the constriction phase. This results in a 9.4 sec full oscillation period, or the frequency of about 0.1 Hz. These results show how a delayed myogenic response can generate oscillations in the PA diameter.

**Fig 6.**
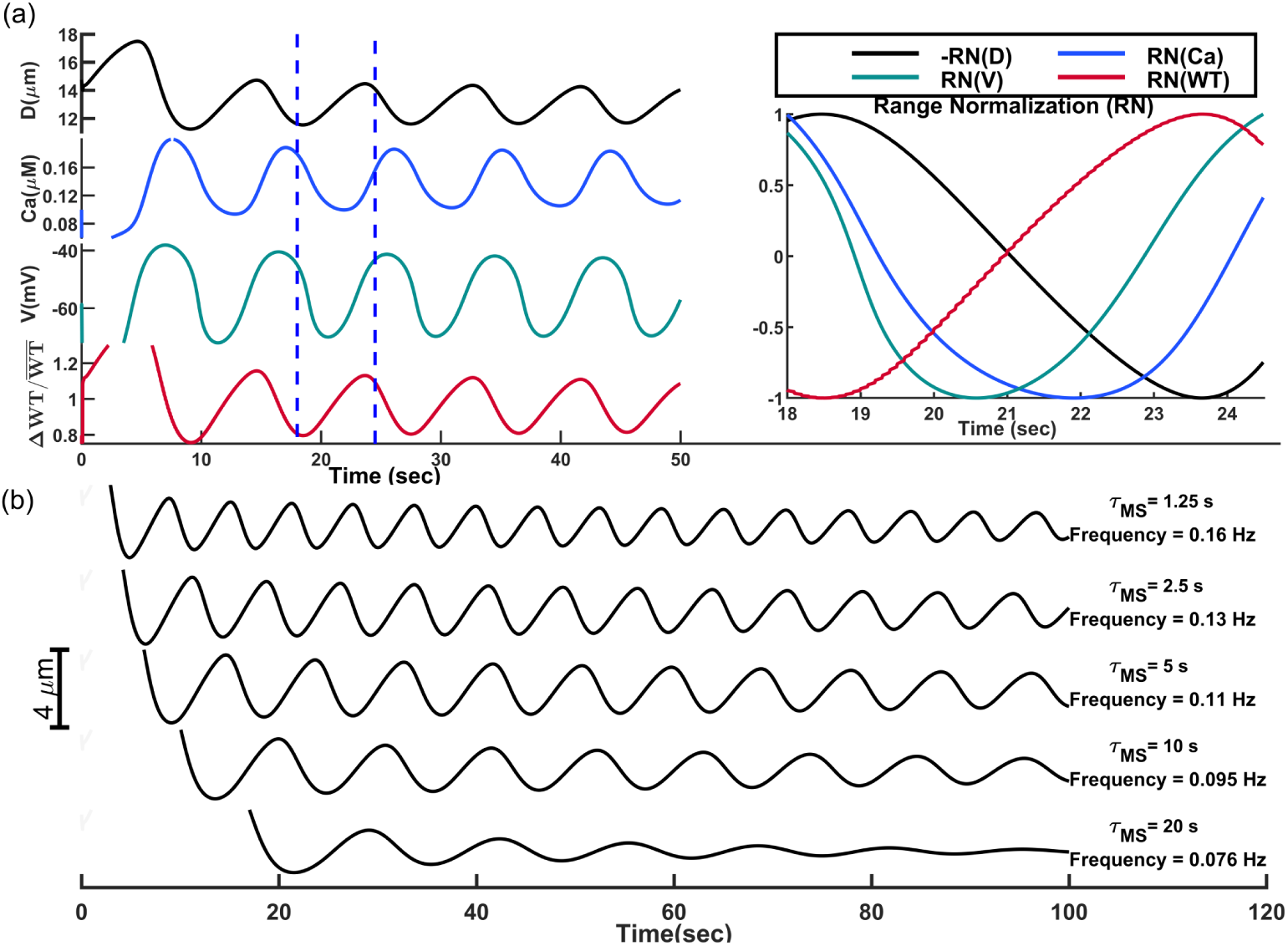
Oscillations of a PA diameter induced by the delayed myogenic responses. (a) Left: Oscillations of the PA segment diameter, SMC [Ca^2+^] and membrane potential, and relative changes of WT. Right: Range-normalized view of oscillations between 18-24.5 sec (the period between blue dashed area in the left panel). The diameter curve is inverted for better visualization of time lags. (b) Analysis of oscillation frequency and amplitude for various mechanotransduction time constants (*τ_mct_*).

Vasomotion oscillations in a PA segment were analyzed for various *τ_mct_* values (Fig. 6(b)). As expected, decreasing *τ_mct_* via increasing NPY concentration (facilitation of the myogenic response) reduces the interval between the maximum WT and the peak of constriction force, and increases oscillation frequency, consistent with experimental observations [105]. In this case, the oscillation amplitude is reduced since the system rapidly corrects any deviations from the target state. The amplitude of vasomotion observed for *τ_mct_*=1.25 sec is smaller than the amplitude of oscillations when *τ_mct_*=5 sec (Fig. 6(b)).

Arteriolar oscillations in the PA model ultimately dampen over time. Under fast myogenic response conditions (small *τ_mct_*), these oscillations fully dampen after 2–3 minutes. When the response is slower (e.g., *τ_mct_*=20 sec), the damping occurs even sooner, within the first minute. Sustained oscillations are observed in the cerebral vasculature in-vivo, indicating that additional factors contribute to maintaining oscillations. To show how these vessels generate synchronized undamped oscillations, we should model PAs as a network of coupled oscillators. In a system of coupled oscillators, synchronized oscillations in adjacent oscillators (PAs) can reinforce each other, sustaining the overall oscillatory activity. For computational efficiency, our simulations so far have modeled only a single PA using cellular-based coupled segments, while keeping all other segments in the cerebrovascular network at a fixed diameter. As a result, the stabilization of proximal hemodynamics led to a gradual damping of oscillations in the analyzed PA. This damping occurs because sustained vasomotion relies on synchronized fluctuations in hemodynamics across the vascular network, which requires coupling between arteriolar segments throughout the system. To simulate this, we developed a computationally efficient non-cellular simplified macro-scale model for all arteriolar segments based on insights gained from the cellular PA model. Then, we used the new simplified model to investigate the effect of network synchronization in sustaining vasomotions.

In the next set of simulations, we initially assigned an ABNP of 60 mmHg to the cerebrovasculature model and adjusted the diameters of all segments to achieve cerebral autoregulation at this ABNP value ([31]). Based on the IP distribution in the network and the assigned diameters, the initial WT in arteriolar segments (AS) was calculated using:

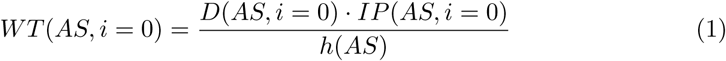

These values, which serve as the baseline around which the WT in arteriolar segments oscillates under an ABNP of 60 mmHg in the cerebrovasculature model, were stored as the *WT_avg_*(*AS*) vector, assuming a constant vessel thickness *h*(*AS*). We further assumed that at this ABNP level, the MT is equal across all arteriolar segments, which results in a hypothetical SMC membrane potential of −45 mV (*V* (*AS, i* = 0) = −45) in all segments. With these initial values set, the simulation loop was initiated with a time step of *dt* = 0.2 sec, where *i* denotes the iteration number. In each iteration, the diameter of each segment was updated according to the macro-scale arteriolar segment model described below. A hemodynamic analysis was then conducted to calculate the new hemodynamics based on updated vessel diameters for the subsequent step. This recursive process was repeated for 600 seconds.

To model vasodynamics in this approach, diameters of arteriolar segments were updated in each iteration based on the delayed myogenic response, passive distension, and the coupling between adjacent segments. Based on the results of Fig. 6(a), if the WT of a segment exceeds *WT_avg_*, the myogenic response is triggered to correct this deviation, albeit with a delay. Consequently, the real-time SMC membrane potential of an arteriolar segment is proportional to the difference between its WT from several seconds earlier and the segment’s *WT_avg_*:

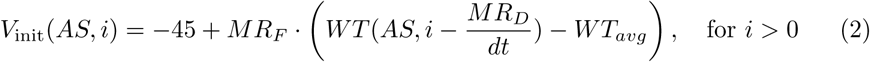

Here, *V*_init_(*AS, i*) represents the initial membrane potential of each arteriolar segment before coupling to adjacent segments. *MR_F_* is the myogenic response factor, and *MR_D_*denotes the myogenic response delay, which was set to 5 sec in our model. As shown in Fig. 6(a), part of the delay in the myogenic response is attributed to the lag between changes in SMC membrane potential and [Ca^2+^], as well as between changes in [Ca^2+^] and adjustments in SMC constriction force. However, in this macro-scale model, where a vessel diameter is adjusted based on the real-time SMC membrane potential, all these delays were encapsulated in *MR_D_*. This means that from the moment WT exceeds *WT_avg_*, until the vessel stops dilating, there is a delay of *MR_D_* sec.

The SMC membrane potential of each segment is influenced not only by its own hemodynamics but also by coupling with adjacent segments. In reality, this coupling occurs through gap junctions, where currents flow based on membrane potential differences, similar to how voltage distributes across resistive-capacitive (RC) circuits. To capture the gradual influence of neighboring segments’ membrane potentials on each other and simulate real-time RC interactions via electrical coupling in our macro-scale model, we implemented a secondary iterative loop within each simulation step. In each iteration of the secondary loop, we computed the gap junction current (*I*_gap_) for each arteriolar segment, progressively updating the segment’s membrane potential in an RC circuit-like manner. At the end of the secondary loop, the final stabilized coupled membrane potential (*V*_final_(*AS, i*)) was obtained from the initial uncoupled membrane potential (*V*_init_(*AS, i*)). The myogenic response (MR) portion of the segment’s diameter was then determined by comparing the segment’s membrane potential to −45 mV:

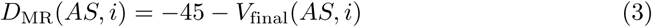

If the final membrane potential of a segment is larger than −45 mV, this means that the WT in that segment (and in surrounding segments due to the coupling) 5 sec ago was above its average value. Consequently, the MR portion of the segment’s diameter becomes negative, leading to a decrease in diameter to counteract the increased WT and bring it closer to the average. Conversely, if the final membrane potential is less than −45 mV, the opposite process occurs.

The passive distension (PD) portion of the segment’s diameter was calculated based on the real-time WT experienced by each segment using the following equation:

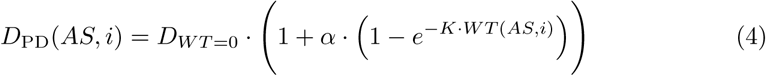

In this equation, *K* represents the rate at which the vessel distends as WT increases, and the parameter *α* determines the maximum magnitude of passive vessel distension achievable at large WT values. The diameter of each segment at each main iteration step was then calculated by adding the passive distension portion of the segment’s diameter (*D*_PD_) and the myogenic response portion of the segment’s diameter (*D*_MR_):

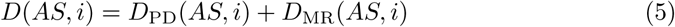

The algorithm corresponding to this simplified model and the model parameter values are provided in the Methods section. Fig. 7(a) shows the oscillatory dynamics of segments at the same cortical depth across 10 different PAs within the simulated vasculature. The oscillations have a square-wave pattern resulting from the vessels alternating between phases of distension and constriction. After the initial passive distension, the vessels remain distended for a fixed period because the initiation of potentiation of MT is delayed. Once this delay elapses, MT increases instantaneously leading to rapid constriction. As WT decreases and crosses below the average threshold, there is another delay before the MT inhibition begins. During this delay, vessels remain constricted despite the small WT. After this delay, MT decreases instantaneously, causing the vessels to dilate quickly. These abrupt transitions, combined with the fixed delays before MT adjustments, produce the distinct square-wave vasodynamics observed.

**Fig 7.**
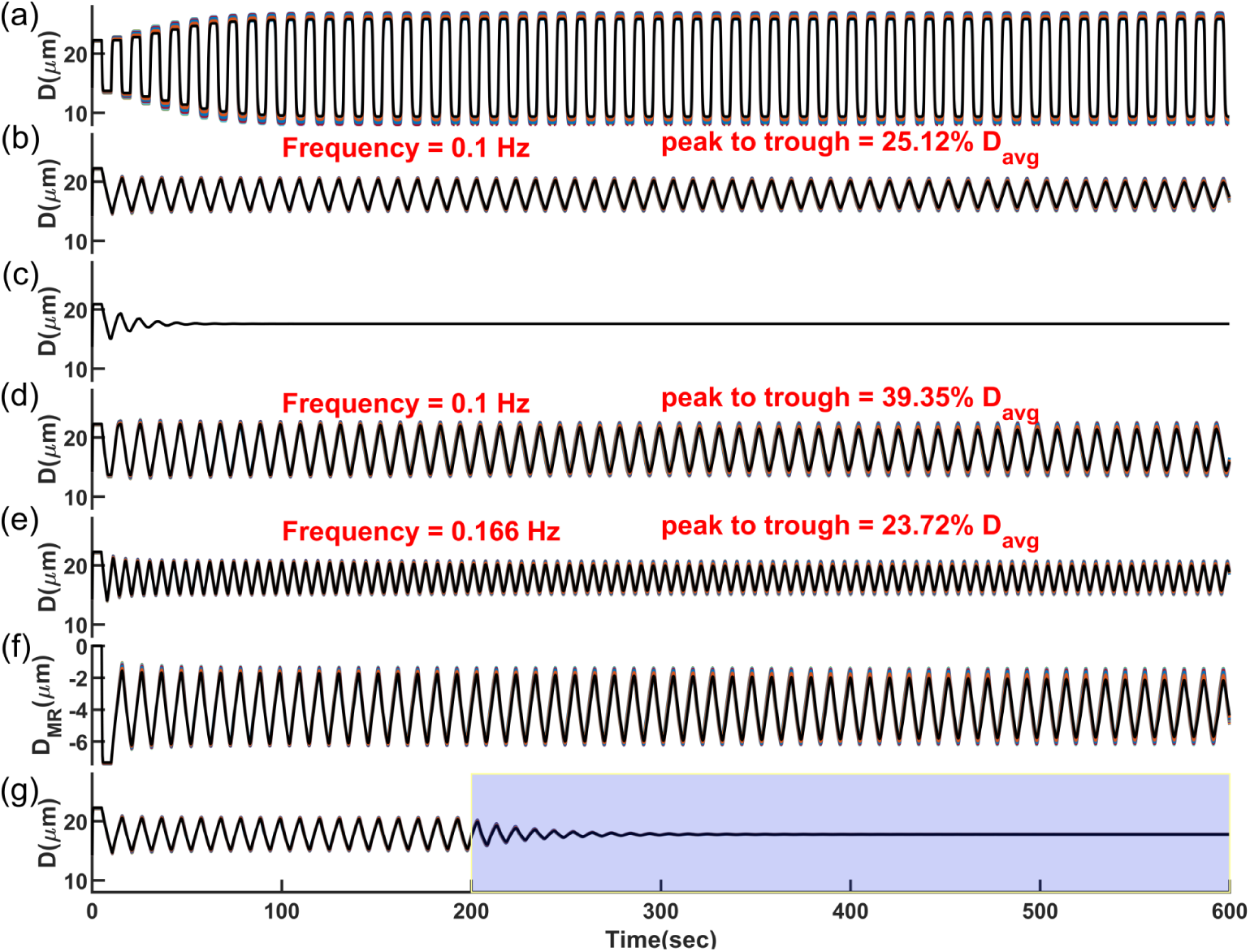
Synchronized vasomotion in a network of 1009 coupled arteriolar segments of the cerebrovascular model, where the diameter of each segment was modeled based on delayed myogenic response (*MR_D_* = 5 s) and passive distension. Curves represent synchronized vasomotion in 10 arteriolar segments with approximately equal diameters from different PAs, overlaid. (a) Square-wave oscillatory pattern with a large amplitude, assuming no constraints on the arteriolar diameter change rate in the model. (b) Triangular and symmetric oscillatory pattern with an amplitude of 25% of the mean diameter, assuming different maximum allowable diameter change rates during dilation and constriction phases (15% per sec for dilation and 7.5% per sec for constriction). (c) Dampened oscillations when the macro-model was implemented only in segments of one PA, rather than in a coupled PA network through pial arteries. (d) Triangular and symmetric oscillatory pattern with an amplitude of 40% of the mean diameter, assuming equal and large maximum allowable diameter change rates during dilation and constriction (15% per sec). (e) Maximum diameter rate change and the delay in the myogenic response are the primary variables modulating vasomotion amplitude and frequency. Larger allowable diameter change rate with a faster myogenic response (*D_MR_* = 3 s) results in a moderate oscillation amplitude (24% of the mean diameter). (f) Myogenic response portion of diameter oscillations in (b). These are negative values as the myogenic response induces constriction to counter passive distension, with constriction periodically potentiated and inhibited. Comparing peak-to-trough diameter oscillations with myogenic response oscillations shows that about 15% of oscillation amplitude is attributed to passive distension. (g) Passive distension plays a key role in sustaining vasomotion. Keeping the passive distension portion of the diameter constant from *t* = 200 sec (highlighted interval) results in dampened oscillations.

In reality, however, MT does not change instantaneously since SMC membrane potential, [Ca^2+^], and the ratio of phosphorylated MLC molecules change slowly over time (see Fig. 6(a)). The fastest observed rate of arteriolar dilation during vasomotion in awake mouse PAs is approximately 10–15% per second ([67]), while the fastest constriction rate was reported to be half of that [106]. To replicate this physiological behavior in our model, we implemented constraints that limit the rate of diameter change for each vessel segment during both dilation (15% per sec) and constriction (7.5% per sec) (Algorithm 2 in the Methods section).

Fig. 7(b) displays the oscillatory behavior of PAs after applying constraints to the diameter change rate. As shown, all PAs synchronously oscillate in a triangular pattern, indicating that synchronization is a dominant feature in these oscillations. This contrasts with the sinusoidal and dampened oscillations observed in the cellular-based PA model depicted in Fig. 6(a). As previously mentioned, when diameters of pial arteries and other PA segments are held constant in the cerebrovasculature model, the hemodynamics in the pial network stabilize, leading to damping of oscillations in the simulated PA. Similarly, Fig. 7(c) shows that implementing the macro-scale model in only one PA within the cerebrovascular model also results in dampened oscillations. These findings show that for the oscillations to sustain, synchronized hemodynamic fluctuations and synchronized changes in SMC membrane potentials are required. These synchronized activities reinforce the oscillations throughout the network.

As depicted in Fig. 7(b), even though the maximum allowable diameter change rates during the dilation and constriction phases were set differently, the vasomotion still has a symmetric triangular pattern. The maximum diameter change rate does not exceed the lower limit of 7.5% per sec. This occurs because the rate of constriction or dilation governs how quickly WT decreases or increases. Consequently, the inhibition or potentiation of the MR also follows the slower rate of constriction or dilation. This prevents any exceedance of the 7.5% per sec limit, even though the model allows for a faster dilation rate. Therefore, the smaller rate of vessel constriction or dilation determines the rate of both phases, leading to a symmetric triangular vasomotion pattern in synchronized vessels. Under this 7.5% per sec constriction rate, the peak-to-trough vasomotion amplitude is approximately 25% of *D_avg_* in our model.

Fig. 7(d) shows that when the maximum allowable diameter change rates were set equal and large in both dilation and constriction phases (15% per sec), the resulting vasomotion amplitude is larger, with a peak-to-trough change of approximately 40% of *D_avg_*. Faster diameter change rates allow vessels to overshoot or undershoot more sharply during the delay period, leading to larger oscillation amplitudes. However, in the physiological vascular system, an increase in the diameter change rate may be accompanied by a decrease in the MR delay, unlike in our model where these variables are independent. This is because faster diameter changes imply quicker changes in SMC [Ca^2+^] and MLC phosphorylation, which are processes that also influence the MR delay. A reduced MR delay limits the vasomotion amplitude, as quicker feedback reduces the time available for energy buildup and overshoot within each cycle (Fig. 6(b)). Fig. 7(e) displays vasomotion with an allowable maximum diameter change rate of 15% per sec for both dilation and constriction, while the MR delay was set to 3 sec. As shown, the amplitude of vasomotion is approximately 24% of *D_avg_*. These simulations show that increasing the vessel diameter change rate, when accompanied by a decrease in the MR delay, and vice versa, have opposing effects on the vasomotion amplitude. This balance helps maintain the vasomotion amplitude within a desired range to prevent excessive vessel constriction or dilation.

While we showed that hemodynamic-vasodynamic interactions can lead to sustained oscillations, we did not explicitly point out the inherent positive feedback process in this dynamic. Delayed negative feedback can initiate oscillations, but these oscillations may not persist indefinitely due to inherent damping mechanisms and energy losses in real physiological systems. Positive feedback mechanisms are essential to amplify oscillations and compensate for these losses. For example, in previous vasomotion models, processes like calcium-induced calcium release (CICR) within SMCs were identified as necessary positive feedback mechanism for sustained oscillations [102, 107]. In our vasomotion model, we propose that passive distension serves as the positive feedback mechanism and it is essential for sustaining vasomotion oscillations. When SMCs relax, the increase in WT leads to further distension of the vessel, creating a positive feedback loop that amplifies dilation. Conversely, when SMCs constrict, the decrease in WT reduces passive distension, amplifying constriction.

Fig. 7(f) shows the MR portion of diameter changes associated with the vasomotion displayed in Fig. 7(b). While the total oscillation amplitude in a vessel segment with an average diameter of about 17.5 *µ*m is approximately 4.5 *µ*m, the MR portion accounts for 3.85 µm (85%), indicating that 0.65 *µ*m (15%) of the oscillation amplitude is attributed to passive distension in the model. Fig. 7(g) displays the vasomotion pattern for *t >* 200 sec when the passive distension portion of vessel diameter was held constant at its value at *t* = 200 sec. As shown, oscillations driven solely by delayed negative feedback start damping immediately. During the first dilation phase for *t >*= 200 sec, the absence of passive distension results in less pronounced dilation, leading to a weaker subsequent constriction as the vessel experienced a smaller maximum WT. This dampening dynamic continues until the vessel diameter stabilizes completely. Collectively, the simulation results in Fig. 7 demonstrate that synchronized hemodynamic-vasodynamic interactions in coupled PAs inherently involve an interplay between delayed negative and positive feedback loops, which serve as the primary drivers of sustained vasomotion.

To provide in-vivo support for the proposed vasomotion model, we used laser speckle contrast imaging (LSCI) to analyze flowmotion in the somatosensory cortex of anesthetized mice (Fig. 8(a)). Flowmotion refers to oscillations in blood flow that originate from vasomotion and have similar characteristics. Five minutes of high-temporal-resolution LSCI signals were recorded from different animals. Fig. 8(b) shows a color-coded LSCI signal in the highlighted region of interest (ROI) for one cycle of flowmotion, which was then spatially averaged across the entire ROI to extract the flowmotion signal, including examples from different animals (Fig. 8(c–e)). In our in-vivo experiments, we rarely observed flowmotion with a complete triangular pattern (Fig. 8(c, d)). This pattern is the characteristic of synchronized, vascular-centric vasomotion generation with minimal interference from neurogenic or astrocytic responses—or phase-locked oscillations of neurogenic, astrocytic, and myogenic activity [17, 108]. These dynamics suggest that, at the time of recording, the animal was deeply anesthetized, and NO production suppressed [109]. NO dampens the myogenic response, and its suppression facilitates the interaction between passive and myogenic responses required for oscillation generation. This facilitated myogenic response may also explain the 0.18–0.2 Hz vasomotion frequency observed in Fig. 8(c), approximately twice the commonly reported 0.1 Hz vasomotion frequency in awake mice [23, 110].

**Fig 8.**
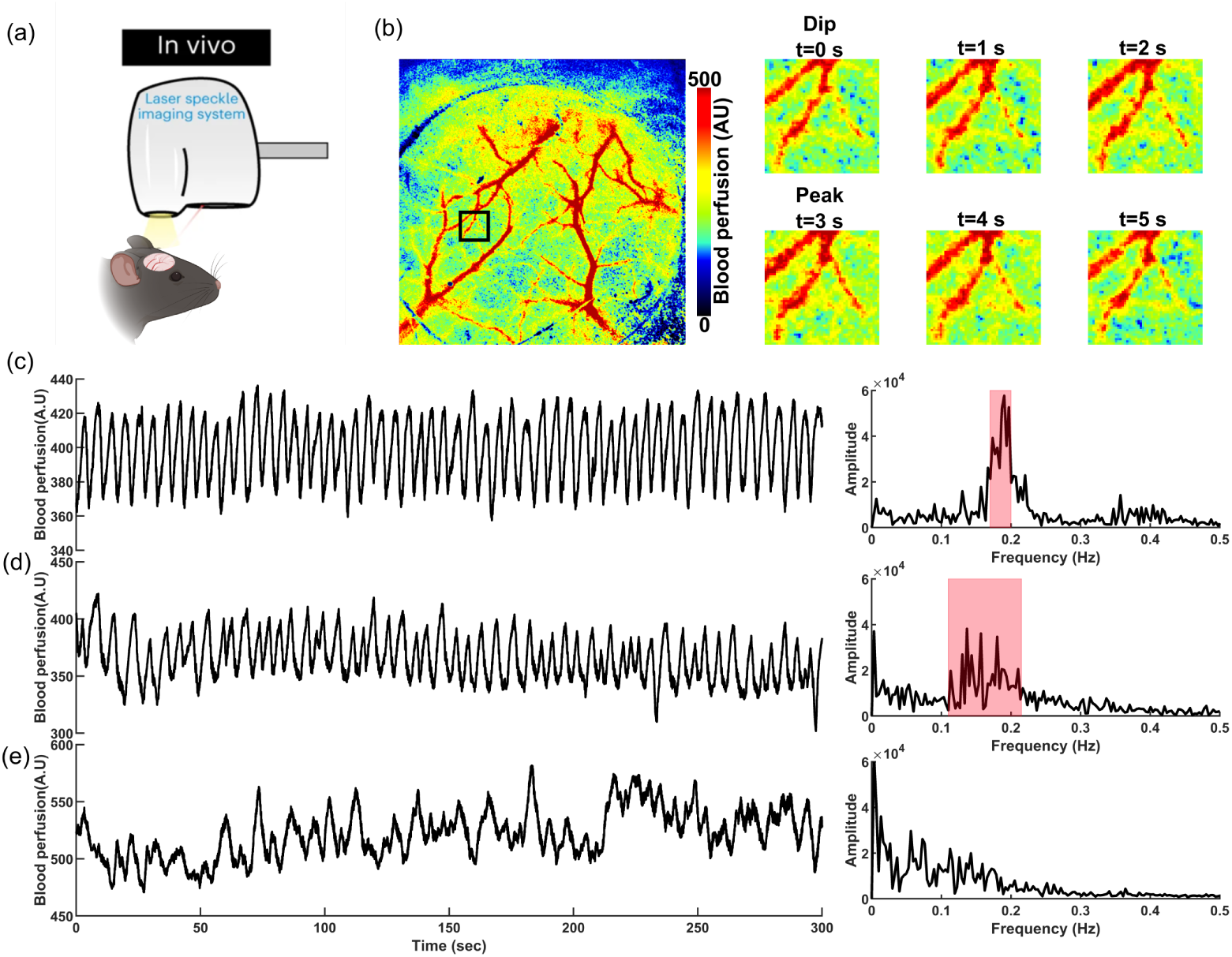
In-vivo analysis of flowmotion characteristics in ketamine-dexmedetomidine anesthetized mice. (a) Schematic representation of in-vivo flowmotion measurements using LSCI. (b) Left: An example of a recorded LSCI signal at a specific time near the left middle cerebral artery (MCA) in the somatosensory cortex of a mouse. Right: Spatiotemporal changes in the LSCI signal during one cycle of flowmotion within the ROI depicted on the left. (c, d) Two examples of LSCI signals where flowmotion exhibits a triangular oscillatory pattern. Even in such triangular flowmotion, there is no dominant oscillatory frequency. (e) A typical recorded flowmotion signal.

Several factors within the mouse brain vasculature can continuously modulate the myogenic response dynamics, including SMC mechantransduction time constant, intracellular Ca^2+^ dynamics, and the sensitivity of MLC phosphorylation to [Ca^2+^]. These factors collectively determine the diameter change rate and MR delay, which can continuously alter the frequency and amplitude of vasomotion (Fig. 7). As shown in Fig. 8(d), within five minutes of recording, multiple dominant frequencies of oscillations emerged in the flowmotion signal. This suggests that, in the dynamic environment of the cerebral vasculature, we should not expect a single dominant frequency arising from an unchanged delayed myogenic response in the network. In our in-vivo experiments, we rarely observed full oscillations, and flowdynamics were generally stochastic (Fig. 8(e)). To investigate the primary drivers of these stochastic flowdynamics, we introduce additional layers to the model.

### Section 3: Modeling NGVC-Driven Arteriolar Vasodynamics

In this section, we first simulate the interplay between passive, myogenic, and neurogenic responses by implementing the proposed NVC model and validate its predictions against experimental data. Subsequently, we examine the role of astrocytes in modulating arteriolar vasodynamics by comparing NVC model-predicted vasodynamics with in-vivo recordings from the awake mouse brain.

#### Modeling Arteriolar Vasodynamics in Response to Neuronal Impulse Activity

To refine the proposed computational model to better reflect physiology, we incorporated in-vivo experimental data.The essential experimental data for our analysis in this section were sourced from in-vivo vasodynamic recordings derived from 0.5 sec optogenetic (OG) stimulation, captured using two-photon microscopy from the somatosensory cortex of mice [111]. Previous studies have shown that OG-evoked vasodilations predominantly initiate below cortical layer IV [112–114]. Thus, analyzing the upward propagation of OG-evoked vasodynamics from deeper to more superficial cortical layers provides a valuable lens to examine the impact of neuronal impulse input on vasodynamics across different cortical depths. This approach allows us to identify depth-dependent deterministic features in neurogenic-myogenic interactions, which can be validated against the predictions of our proposed model to evaluate its accuracy.

Brief OG stimulation triggers delayed and attenuated arteriolar dilations in superficial cortical layers compared to deeper ones, with a more pronounced post-stimulus undershoot in the deeper layers ([112, 115]) (Fig. 9(a,b)). We quantified the propagation speed of arteriolar dilation by measuring the onset and peak times of dilation at different cortical depths of each PA (as detailed in Methods), and identified vasodilation propagation speeds of 300-400 *µ*m/sec (Fig. 7(c)), significantly slower than the retrograde electrical propagation speed of 2 mm/sec reported by [10].

**Fig 9.**
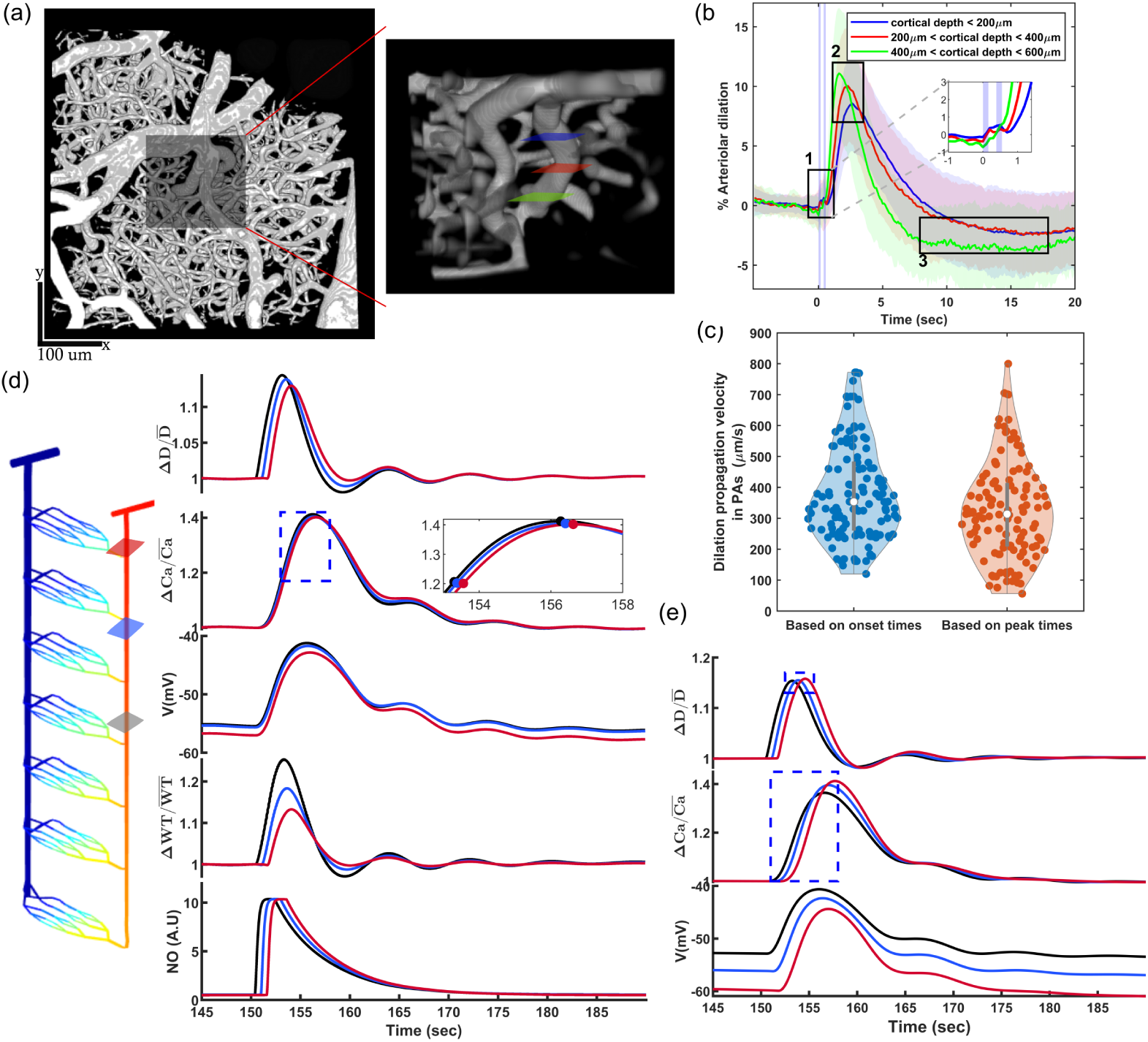
Neurogenic impulse response of the CBF regulatory system: (a-c) OG-evoked vascular reactivity in PAs of awake mouse brain across different cortical depths, sourced from [111, 112]. (a-left) Example of a two-photon image of the mouse somatosensory cortex vasculature. (a-right) Depth-resolved illustration of the ROI shown on the left. (b) Averaged OG-evoked arteriolar vasodynamics measured in three different cortical depth regions (colored cross-sections in (a-right)), highlighting three depth-dependent features of the in-vivo OG-evoked vasodynamics: 1. Delayed arteriolar vasodilation in superficial layers, 2. Attenuated arteriolar dilation in superficial layers, and 3. More pronounced post-stimulus constrictions in deeper layers. (c) Violin plot of the calculated speed of vasodilation propagation along each PA in the dataset, determined using onset-to-onset and peak-to-peak times from concurrently recorded vasodynamics across different depths, assuming PAs penetrate perpendicularly into the tissue. (d) Left: Modeled PA consisting of 28 coupled segments, each including EC, SMC, and passive distension models. Right: Spatiotemporal changes in model parameters at three cross-sections of the PA, located at depths of 405, 225, and 45 *µ*m, in response to instantaneous neuronal activity. The dots in the zoomed-in area represent timestamps used for calculating Ca^2+^ wave propagation and electrical signaling. (e) Spatiotemporal changes in model parameters in response to instantaneous neuronal activity, assuming there exists no electrical coupling between PA segments (removal of gap junctions), resulting in inaccurate predictions of vasodynamics compared to in-vivo recordings.

nNOS-expressing interneurons evoke arterial dilation in the somatosensory cortex of mice [116], and the delayed vasodilation in superficial arterioles might result from slower neurovascular communication kinetics in superficial layers rather than from the delayed propagation of neural activity from the deeper layers to the superficial area [112, 115]. Taking these observations together, we assumed that instantaneous neuronal activity leads to the release of equal amount of neuronally produced NO across all cortical depths, with delayed modulation of the myogenic response and vasodilation in superficial layers due to slower neurovascular communication kinetics. We tested whether this scenario could account for the depth-dependent deterministic features observed in PA vasodynamics, including the attenuated vasodilation during upward propagation and the more pronounced post-stimulus undershoot in deeper layers.

We initially set the ABNP of the cerebrovascular model to 60 mmHg, and we allowed the oscillations to fully damp before applying neurogenic input to the cellular-level PA, allowing us to analyze the average neurogenic impulse response of the PA, without interference from oscillations. We then assumed that instantaneous neuronal activity induced a sudden rise in NO concentration in PA segments located below 600 *µ*m depth, whereas in PA segments above 600 *µ*m, this rise propagated at a speed of 400 *µ*m/sec. This delayed adjustment was designed to capture the depth-dependent onset of vasodilation observed in Fig. 9(b).

Fig. 9(d) shows the changes in various parameters of the analyzed PA at three different depths. As shown in the bottom panel, the NO/cGMP concentrations were modeled to increase with a delay corresponding to cortical depth and to decay exponentially with a 5 sec time constant. The NO/cGMP/PKG pathway primarily induces rapid vasodilation by activating PKG, which in turn activates MLCP and desensitizes MLC to Ca^2+^ which leads to partial dampening of the myogenic response (Section 1). Therefore, when the PA segment located at 405 *µ*m (black curves) experiences this NO-induced vasodilation, the damped myogenic response is followed by an increase in vessel segment diameter and elevated IP during the dilation phase, which increases the WT and potentiates the myogenic response. This, in turn, causes a subsequent increase in SMC membrane potential and SMC [Ca^2+^]. Due to the electrical coupling between vascular cells, the increased SMC membrane potential in deeper PA segments, which experience the rise in NO concentration earlier, is transmitted to the upper vascular cells via gap junctions, causing rapid depolarization in the upper layers’ SMCs even before the NO concentration rises there. Based on the timing of the mid and peak [Ca^2+^] increases in analyzed segments, the electrical propagation speed in our model was approximately 1200 *µ*m/s. In-vivo findings showed that during instantaneous sensory stimulation, the propagation of vasodilation in superficial vessels lagged behind the rise in SMC [Ca^2+^], indicating that vasodilation occurred at a speed of around 400 *µ*m/sec, consistent with the values calculated in Fig. 9(c), where the SMC Ca^2+^ wave was propagating at approximately 1200 *µ*m/sec [117].

During the time interval when SMC [Ca^2+^] is increasing almost simultaneously across all segments but before reaching the peak, the NO/cGMP concentration is higher in the deeper layers due to the earlier rise. As a result, the dilation of deeper PA segments during this phase is more pronounced than in the superficial segments. In the superficial segments, the equal concentration of NO/cGMP is accompanied by a more potentiated myogenic response evoked by earlier dilations in deeper segments, which is quickly transmitted electrically upstream (Fig. 9(d) - top panel). This dynamic explains the attenuation observed in the upward propagation of vasodilation observed experimentally (Fig. 9(b)). Furthermore, the potentiated myogenic response results in rapid vasoconstriction following the initial NO-induced partial dampening of the myogenic response, rather than the vasodynamic response that follows the slow degradation of NO/cGMP. Therefore, during the interval when SMC [Ca^2+^] is near its peak in all segments, the NO/cGMP concentration in deeper cortical layers is lower due to its earlier degradation following the initial rise. At this interval, the increased MLC Ca^2+^ sensitivity in the deeper segments leads to a more pronounced post-stimulus undershoot. This dynamic also explains the more pronounced post-stimulation undershoot observed in the deeper segments experimentally (Fig. 9(b)).

Fig. 9(e) shows the simulation results under similar neurogenic input, but without coupling between adjacent vascular cells in the modeled PA. The basal SMC membrane potential at different depths is not within the same range as it was in the coupled segments scenario depicted in Fig. 9(d). Additionally, the myogenic response begins to potentiate independently in each PA segment following its delayed NO-induced vasodilation, rather than in a synchronized manner, despite the delay in dilation. In this simulation, the depth-dependent deterministic features observed experimentally and computationally are absent. Therefore, Fig. 9 collectively demonstrates that during instantaneous neuronal activity, neuronally released NO acts on SMCs to momentarily dampen the myogenic response and induce vasodilation. This NO-induced dilation subsequently potentiates the myogenic response, leading to the attenuation of the initial vasodilation and the emergence of a post-stimulus undershoot in PA segments. Depending on the cortical depth of the PA, the degree of attenuation in the initial vasodilation and the magnitude of the post-stimulus undershoot vary.

Next, we analyzed the effects of different *τ_mct_* values on vasodynamics in response to neuronal impulse input. Increasing *τ_mct_*—which corresponds to lower activation of SMCs’ NPYRs—resulted in a slower myogenic response, where adjustments allowed a less immediate and pronounced reaction to changes in the diameter and WT (Fig. 10(a)). Consequently, the initial dilation induced by NO-dependent partial dampening of the myogenic response was smallest with the fastest myogenic response and largest with the slowest. However, the magnitude of the post-stimulus undershoot depends not only on the degree of the potentiated myogenic response, which is more pronounced with smaller *τ_mct_*, but also on the degradation rate of NO/cGMP molecules. The post-stimulus undershoot was most pronounced at *τ_mct_* values of 25 and 50 secs, compared to both smaller and larger *τ_mct_* values. The reason is, within this *τ_mct_* range, and with a 5 sec time constant for the exponential decay of NO/cGMP concentration, the increased Ca^2+^ sensitivity of MLC from 10 to 15 sec after the initial NO/cGMP rise coincides with a delayed myogenic response-induced Ca^2+^ peak. This, in turn, resulted in a more pronounced post-stimulus undershoot compared to faster myogenic response scenarios, where even larger Ca^2+^ peaks occurred within the first 10 sec, when MLC sensitivity to Ca^2+^ was reduced due to higher NO/cGMP concentration. Furthermore, the analyzed dynamics shows that both the duration of the initial vasodilation and the post-stimulus undershoot increase with increasing *τ_mct_*, indicating a slower myogenic response. These modeled and comparative dynamics were similarly observed in instantaneous sensory-evoked vasodynamics in adult and aged mice [118]. Based on the reported vasodynamics and concurrent SMC Ca^2+^ monitoring in that study, and the vasodynamics we analyzed under various *τ_mct_* values in Fig. 10(a), we can conclude that the myogenic response in aged mice is slower than in adults, since the duration of both vasodilation and the myogenic response-induced elevation of SMC [Ca^2+^] were longer in aged mice.

**Fig 10.**
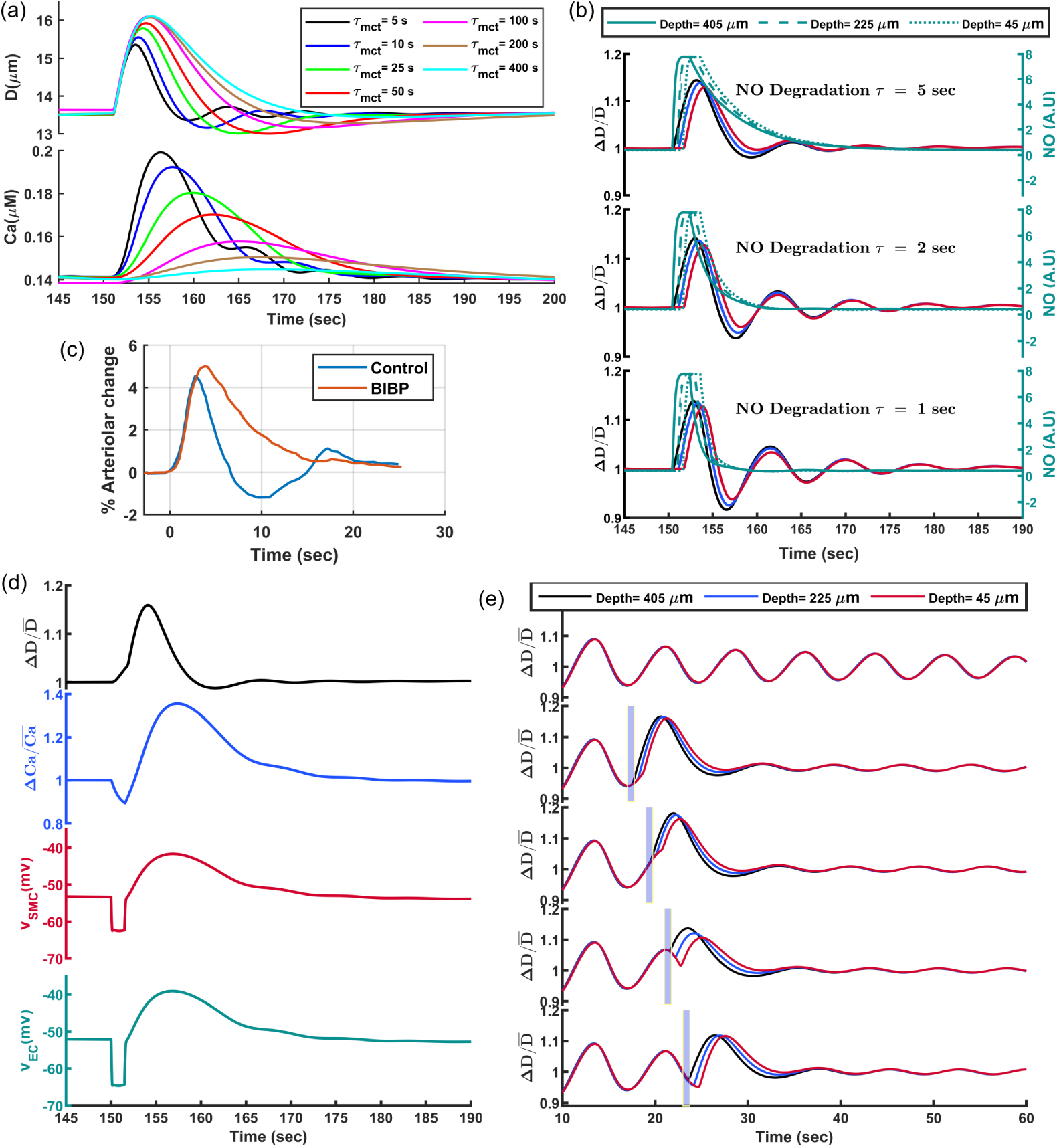
Analyzing arteriolar vasodynamics in response to neuronal impulse input under various model parameter values. (a) Vasodynamics and SMC Ca^2+^ dynamics under various SMC mechanotransduction time constant (*τ_mct_*). Decreasing *τ_mct_* leads to a more attenuated vasodilation, along with a shorter duration of vasodilation and post-stimulus undershoot. (b) Predicted vasodynamics under various NO/cGMP degradation time constants. Faster degradation results in more pronounced post-stimulus constriction and oscillatory behavior, which is absent in in-vivo signals, suggesting that NO/cGMP degradation occurs slowly under physiological conditions. (c) In-vivo recorded vasodynamics from Uhlirova’s study [112], showing significant changes in vasodynamics upon the application of the Y1 receptor blocker BIBP 3226. Data were digitized from Uhlirova’s study and replotted here, under the terms of the CC BY 4.0 license (https://creativecommons.org/licenses/by/4.0/). (d) Model-predicted vasodynamics assuming that, during instantaneous neuronal activity, neuronally released mediators also hyperpolarize vascular cells, leading to biphasic SMC [Ca^2+^] and larger vasodilation. (e) Model-predicted vasodynamics based on the assumption that instantaneous neuronal activity occurs at four different phases of ongoing vasomotion.

Next, we analyzed vasodynamics in response to neuronal impulse input with a fixed *τ_mct_* of 5 sec, while changing the time constant of the assumed NO/cGMP concentration exponential decay. As shown in Fig. 10(b), reducing the time constant, which accelerates the decay of NO/cGMP concentration, affected the duration of vasodilation, the degree of post-stimulus undershoot, and the timing of its occurrence. Comparing these simulation results with the experimental data shown in Fig. 9(b) suggests that a slower decay in NO/cGMP concentration, with *τ* = 5 sec degradation time constant, replicates the experimentally observed vasodynamics more accurately. Therefore, the 5-sec time constant is a reasonable estimate for the NO/cGMP concentration exponential decay.

Our systems biology approach demonstrated that the potentiated myogenic response is primarily responsible for the biphasic vascular reactivity observed in neuronally-evoked vasodynamics, characterized by an initial vasodilation followed by rapid constriction, often leading to a post-stimulus undershoot. However, the constriction phase is blunted following the administration of the NPY receptor antagonist, BIBP ([112]), suggesting a potential interaction between NPY-positive inhibitory neurons and vascular SMCs. This interaction would facilitate inhibitory response within the CBF system (myogenic response), effectively counterbalancing initial blood flow increases triggered by excitatory responses. By comparing the vasodynamics under various SMC *τ_mct_*values (Fig. 10(a)) with those observed after BIBP administration (Fig. 10(c)), where neither the attenuation of initial vasodilation nor the post-stimulus undershoot is present, we can infer that the NPY receptor antagonist significantly slows the myogenic response. In our model, to replicate vasodynamics under BIBP application, the SMC *τ_mct_* would have to increase by a factor of 40-80 or more compared to *τ_mct_* = 5 sec. Therefore, the vasodynamics observed with BIBP likely represent pure NO-induced vasodilation, which follows the decay of NO/cGMP concentration. This approximately 16-20 sec constriction phase suggests that the NO/cGMP degradation time constant is around 4-5 sec.

In our analysis of vasodynamics in response to neuronal impulse input, we assumed that the brief duration of neuronal activity was insufficient to induce SMC hyperpolarization, either through neuronally released potassium acting on the endothelial network or by direct neuronal activation of SMC glutamate receptors colocalized with BK channels (Section 1). However, if either pathway is activated, the resultant vasodynamics would not change significantly. Fig. 10(d) shows the vasodynamics where the SMC membrane potential is immediately hyperpolarized due to neuronally mediated increases in EC [*K*^+^]_ex_ and the activation of SMC BK channels for a brief period (1.5 sec). This rapid hyperpolarization of SMCs caused an initial decrease in SMC Ca^2+^ levels, followed by a myogenic response-dependent elevation in SMC Ca^2+^, resulting in a biphasic Ca^2+^ pattern—an initial decline followed by a subsequent increase. This biphasic Ca^2+^ pattern has been observed in several in-vivo studies which were analyzing concurrent SMC Ca^2+^ dynamics and vasodynamics ([117, 118]). The initial SMC hyperpolarization led to a slightly more pronounced vasodilation due to cumulative effects of neuronal mediators inhibiting and dampening the myogenic response, but with no notable differences emerging after this initial vasodilation compared to scenarios without SMC hyperpolarization. This response might change under faster Ca^2+^ dynamics in SMCs, where a hyperpolarizing spike in membrane potential would lead to a corresponding rapid decrease in [Ca^2+^].

Next, we analyzed vasodynamics in response to neuronal impulse input, assuming that there is ongoing vasomotion in the arteriolar segments. As previously mentioned, the cellular-level PA model oscillates for two to three minutes after the simulation onset, until oscillations are fully damped (Fig. 10(e) - top panel). Here, we assumed that neuronal input was applied to the PA model at four distinct points in time during oscillations including the: oscillation trough, mid-dilation, peak, and mid-constriction. As shown in Fig. 10(e), the ongoing oscillatory myogenic response influences both the magnitude of the initial dilation and the post-stimulus undershoot of vasodynamics depending on the timing of the neurogenic input to the PA. Furthermore, since NO suppresses the myogenic response for 10-20 sec post-stimulation due to its slow degradation rate, the ongoing vasomotion is damped after the stimulus. This occurs because the vessel is less constricted following the initial stimulus-evoked vasodilation, and the reduced constriction leads to a weaker delayed inhibition of the myogenic response during the dilation phase. As a result, oscillations remain damped and do not recover in this simulation, since we only modeled one PA with no synchronization in the cerebrovascular model. In reality, synchronization in the vascular network may help the oscillation recover post-stimulation, albeit with a new phase compared to the pre-stimulation oscillation, and this new phase is determined by the timing of the stimulus relative to the ongoing oscillation (Fig. 10(e)).

#### Modeling Arteriolar Vasodynamics in Response to Neuronal Step Activity

Here, we analyze vasodynamic output in response to a neuronal step input, assuming a relatively stable overall neural activity and minimal neuronal feedback inhibition during sustained stimulation. If the stimulation persists for an extended duration, the neuronally mediated SMC hyperpolarization, in addition to the dampening effect of NO on the myogenic response, remains effective for the entire period of neuronal activity. Fig. 11(a) shows key PA model variables in response to a 10 sec sustained neurogenic input under three different neuronal activity levels with *τ_mct_* = 5 sec. We assumed that the hypothetical values of all neuronally released mediators, including NO, glutamate, and discharged potassium, increase linearly from level 1 to level 3. As shown, the SMC membrane potential initially hyperpolarizes in proportion to the level of neuronally released SMC hyperpolarization mediators. Simultaneously, NO-induced dilation significantly potentiates the myogenic response, which counteracts the neuronally induced SMC hyperpolarization, causing the SMC [Ca^2+^] to revert toward its pre-stimulation level or even higher after several seconds, consisntant with in-vivo studies [12, 119].

**Fig 11.**
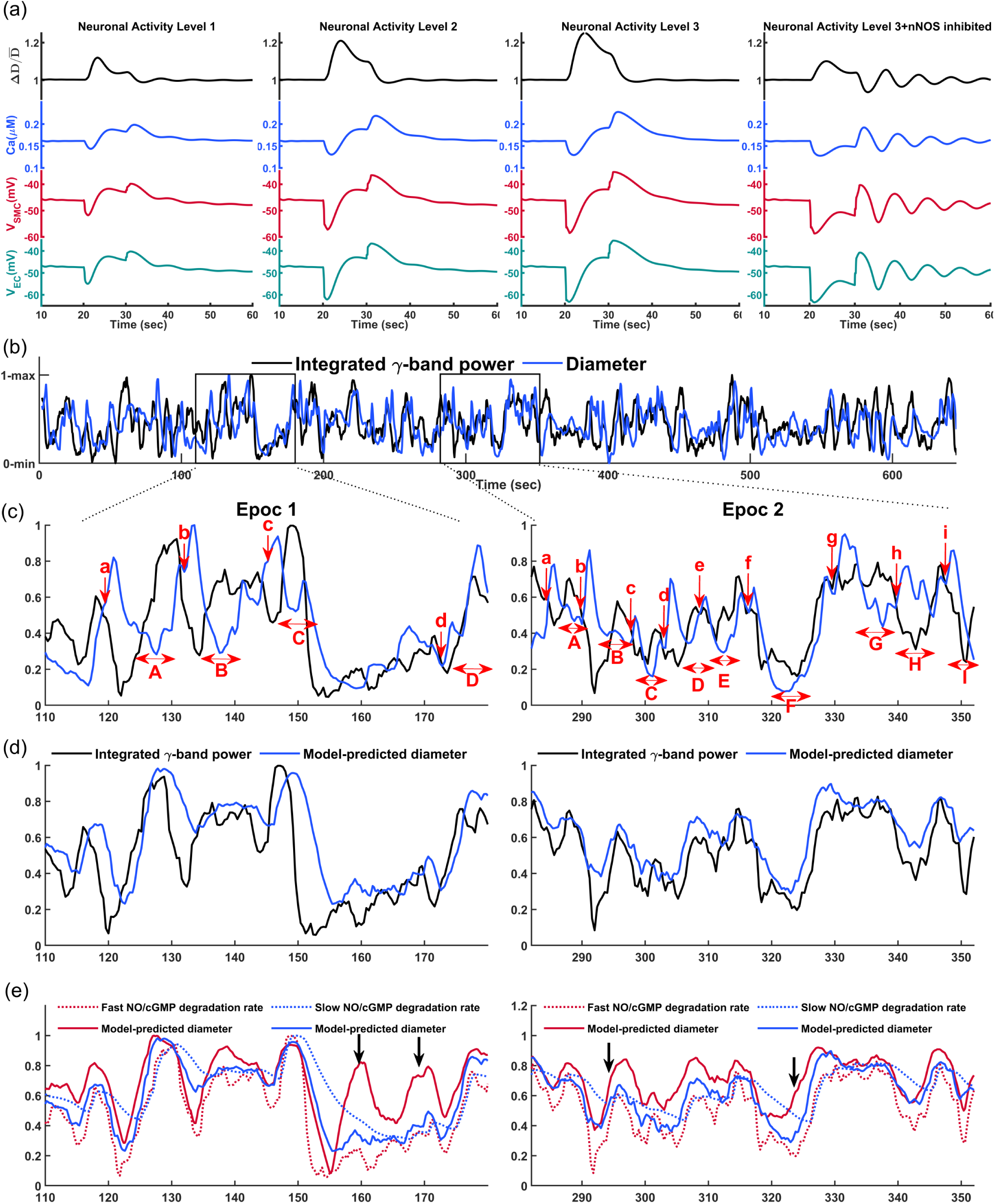
membrane potential, and SMC [Ca^2+^] are plotted for three hypothetical neuronal activity levels, increasing from level 1 to level 3. The corresponding hypothetical level of neuronal mediators are as follows: EC external potassium concentration changes from 3 mM at the baseline to 4.5, 6, and 7.5 mM; Glutamate concentration changes from 0 at baseline to 0.4, 0.8, and 1.2 AU; NO levels change from 0 at baseline to 2.5, 5, and 7.5 AU. In the right panel, all mediators corresponding to level 3 neuronal activity were applied except NO production. (b–e) Role of astrocytes in modulating arteriolar vasodynamics: (b) In-vivo recorded integrated gamma-band power and concurrent vasodynamics of an awake mouse as reported by Mateo et al. [17]. (c) Magnified view of two range-normalized epochs of in-vivo signals. Single-sided arrows and lowercase letters indicate timestamps for the initiation of astrocytic responses triggered by vessel constriction. Double-sided arrows and capital letters indicate time intervals for delayed potentiated myogenic responses in reaction to astrocytic responses. (d) Predicted vasodynamics by the proposed NVC model. (e) Predicted vasodynamics by the NVC model under both slow and fast degradation rates of NO/cGMP.

The myogenic response should get more potentiated in neuronal activity level 3 compared to level 2, as the dose-dependent NO-induced vasodilation in level 3 is more pronounced. Consequently, we would expect a larger myogenic response-induced Ca^2+^ elevation in level 3 compared to level 2, although both reach the same [Ca^2+^] and SMC membrane potential by the end of the stimulation. This occurs because the more pronounced potentiated myogenic response in level 3 is counteracted by a stronger neuronally induced SMC hyperpolarization (inhibition of the myogenic response). As a result, the steady-state myogenic tone at the end of stimulation in both levels 2 and 3 are approximately equal. This dynamic is evident when comparing the level of SMC membrane potential depolarization immediately after the stimulus ends. Upon the sudden cessation of neuronal inputs, the SMC depolarization in level 3 is more than level 2, indicating that neuronally mediated inhibition of the myogenic response during the stimulation period was larger in level 3 than level 2.

Next, we analyzed the vasodynamics in neuronal activity level 3, assuming that nNOS is inhibited and NO is not produced during the stimulation period (Fig. 11(a-right panel). As shown, the neuronally mediated SMC hyperpolarization induced initial dilation which was followed by the myogenic response potentiation and a constriction phase. These simulation results demonstrate that delayed myogenic response potentiation is not solely attributed to NO-induced dilation, and any form of sudden constriction or dilation is followed by the subsequent dilation or constriction, respectively. Post-stimulus, with the cessation of neuronal activity, the SMC depolarizes, and the vessel immediately constricts since there is no NO to desensitize the coupling between changes in Ca^2+^ and the constriction force of SMCs. Subsequently, this immediate constriction is followed by a dilation phase caused by the delayed myogenic response, which is then followed by another constriction phase. Therefore, comparing the vasodynamics in level 3 neuronal activity with and without nNOS inhibition indicates that most of the of vessel dilation during the stimulation period is induced by NO. Moreover, vessel diameter oscillations, post-stimulus, are more pronounced in the absence of NO.

#### Astrocytic Regulation of Arteriolar Vasodynamics

Ultra-slow fluctuations in neuronal signaling, occurring as an envelope over *γ*-band activity, entrain vasomotion [17]. Therefore, we aimed to test whether modulating arteriolar myogenic response with the envelope of *γ*-band activity would result in model-predicted vasodynamics that align with experimental observations. Fig. 11(b) shows an example of recorded integrated gamma-band power over 0.4 sec intervals (the envelope of *γ*-band activity) and concurrent arteriolar vasodynamic measurements are as reported in Mateo et al.’s study. For this test, we used the simplified macro-scale arteriolar segment model, where vessel diameter was determined by delayed myogenic response in coupled segments and passive distension, which resulted in vasomotion in resting state. To include neuronal modulation in this model, we needed to determine the time lag between changes in neuronal activity and the corresponding changes in the myogenic response, whether through inhibition or dampening. Vasodynamics lag behind the envelope of *γ*-band activity by approximately 1.9 sec ([17]). Therefore, we can modify Eq. 3 as follows:

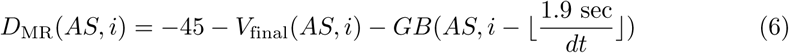

Here, 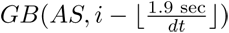 represents a factor of the range-normalized *γ*-band signal, where the real-time myogenic response is inhibited by the *γ*-band signal from 1.9 sec earlier. We showed that the myogenic response is damped by NO produced during neuronal activity, and that NO/cGMP degradation occurs gradually. Therefore, we initially smoothed the decreasing trend of the range-normalized *γ*-band signal and then used this smoothed signal to dampen the myogenic response. Fig. 11(e-blue dashed line) shows the smoothed range-normalized *γ*-band signal, while Fig. 11(e-red dashed line) depicts the non-smoothed version. Thus, the real-time vessel diameter can be updated as:

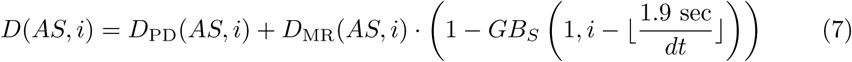

The term 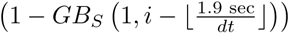 models the dampening effect of neuronally-produced NO on the myogenic response by the smoothed and delayed *γ*-band signal.

Fig. 11(c) displays a magnified view of two range-normalized epochs of in-vivo signals, and Fig. 11(d) shows the range-normalized predicted vasodynamic by our macro-scale NVC model, plotted alongside the in-vivo *γ*-band signal. Comparing the model-predicted vasodynamic with the in-vivo vasodynamic shows t hat, in some cases, the model output shows good similarity to the in-vivo signal. For instance, in epoch 1, between the 145-170 sec interval, the smoothed decay of NO-induced dampening of the myogenic response prevents the immediate resumption of oscillatory myogenic activity after a sudden decrease in neuronal activity. As shown in the red curves in Fig. 11(e), if the NO/cGMP concentration closely follow the *γ*-band signal without smoothing (red dashed curve), the rapid decline in diameter would trigger a pronounced delayed myogenic response, leading to a subsequent dilation and the onset of oscillations. Therefore, the slow decay of NO-induced dampening of myogenic response prevents the emergence of oscillations in arteriolar vasodynamics.

Even with the above refinements, significant discrepancies exist between the vasodynamics predicted by the NVC model and the in-vivo vasodynamics, which are associated with the astrocytic response in the CBF regulatory system. For example, take the timestamp ‘b’ in epoch 1. At this point, our model predicted that, due to the decline in the *γ*-band signal, the vasodynamics would follow a delayed downward trend. However, during the initial moments of this decline, we observed a sudden and unexpected rise. This unexpected deterministic feature is not exclusive to this timestamp; similar events could be seen at all timestamps marked by red single-sided arrows and lowercase letters. For instance, at timestamps ‘a’ and ‘c’ in epoch 1, similar unexpected events occur, and in epoch 2, we observed at least nine similar events within the 70 sec magnified view.

We attributed this deterministic feature of vasodynamics to astrocytes activities based on studies investigating astrocytes contributions to FH. A key study demonstrating that astrocytes possess mechanosensitive TRPV4 channels, which enable them to detect mechanical strain caused by changes in vessel diameter [120]. If, due to ongoing neuronal and synaptic activities, astrocytic endfeet TRPV4 channels are preconditioned with their endogenous ligands, such as epoxyeicosatrienoic acids (EET) [12, 121], and if IP3 receptors on the endoplasmic reticulum within astrocytic endfeet are primed for a Ca^2+^ wave [122], paired with astrocytes depolarization caused by potassium removal from brain tissue, then TRPV4-evoked Ca^2+^ signaling in astrocyte endfeet could activate nearby BK channels. This would trigger the release of retained potassium into the surrounding perivascular space (PVS), amplifying the conductance of SMC Kir channels and resulting in SMC hyperpolarization (Fig. 4(a)). Thus, if astrocyte endfeet are primed for significant Ca^2+^ signaling and subsequently detect vessel constriction during FH, they can actively induce arteriolar vasodilation. This mechanism aligns with findings from multiple studies, where Ca^2+^ signaling in astrocyte endfeet was observed post-stimulation in short-duration FH experiments, specifically when PAs began to constrict [123–127].

At timestamp ‘b’ in epoch 1, after astrocyte endfeet detected vessel constriction and released a large amount of potassium near the SMC Kir channels, the vessel dilated. Following this immediate rise in vessel diameter, the delayed myogenic response induced a subsequent constriction phase. Although our model predicted that vasodynamics should remain relatively unchanged during time interval ‘B’, the delayed potentiated myogenic response triggered by astrocytic activity at timestamp ‘b’ induced a constriction phase, despite neuronal activity remaining relatively high during the ‘B’ interval. Similarly, during time interval ‘C’ in epoch 1, while our proposed NVC model predicted that vasodynamics would follow the rising trend of the *γ*-band signal with a delay, the delayed potentiated myogenic response triggered by astrocytic inhibition of the myogenic response at timestamp ‘c’ had a significant impact on vasodynamics. Once this delayed potentiated myogenic response passed, the increased *γ*-band signal caused vessel dilation, but this dilation was quickly terminated due to the subsequent decline in the *γ*-band signal. At timestamp ‘f’ in epoch 2, while our NVC model predicted constriction, astrocytic inhibition of the myogenic response caused the vessel to dilate. However, neuronal activity also showed an immediate decline during this time. As a result, during time interval ‘F,’ both the delayed potentiated myogenic response triggered by astrocytic activity at timestamp ‘f’ and the small neuronal inhibition of the myogenic response significantly constricted the vessels. Even the subsequent increase in the *γ*-band signal could not easily elevate the vessel diameter during this interval, giving the impression that the *γ*-band signal would lag behind the vasodynamics. However, this was not the case, since the delayed potentiated myogenic response triggered by astrocytic activity drove this unexpected vasodynamics.

Other indicated timestamps for the initiation of astrocytic responses and the corresponding time intervals for delayed potentiated myogenic responses in reaction to these astrocytic signals are well-justified with the proposed NGVC model. As demonstrated, astrocytes periodically inhibit the myogenic response, particularly in response to its potentiation, while neurons continuously send signals to modulate the myogenic response in vessels. Study of these interactions provide a clearer understanding of how astrocytes introduce additional low-frequency components to vasodynamics [120]. Vasodynamics generally should have stochastic low-frequency components from the envelope of the *γ*-band signal, as the myogenic response is continuously inhibited or dampened by this signal. Additionally, the non-singular frequency of vasomotion, which is more pronounced during periods of low and steady *γ*-band signal amplitude, also emerges in vasodynamics. Our findings show that astrocytic activity further modulates arteriolar vasodynamics, thereby adding more low-frequency components to the vasodynamics.

## Discussion

### Significance and summary

In this study, we introduced a novel framework for mouse cerebral arteriolar vasodynamics, incorporating interactions among passive, myogenic, neurogenic, and astrocytic responses of the CBF regulatory system. Under resting conditions, passive distension and delayed myogenic responses generate vasodynamic oscillations; however, during dynamic brain activity, neurogenic and astrocytic inputs modulate the myogenic response, disrupt oscillations, and introduce vasodynamic/hemodynamic fluctuations that are approximately 60% correlated with fluctuation in neuronal activity [17, 128]. We identified three state-dependent elements in the modulation of the myogenic response that explain the vasodynamic fluctuations not directly correlated with neuronal activity (Fig. 11(c)):

1. The modulation of the myogenic response by neuronal and astrocytic inputs is state-dependent, with the degree of modulation determined by the vascular WT seconds before the input is received.
2. The dynamic of myogenic response is state-dependent and governed by the balance between dampening modulators (e.g., NO/cGMP/PKG/MLCP activation) and facilitatory modulators (e.g., NPY receptor activation). When this balance shifts toward a facilitated myogenic response, sudden vasodynamic changes would induce oscillatory vasodynamics that may not be directly correlated with the ongoing neuronal activity.
3. The modulation of the myogenic response by astrocytes is state-dependent and is periodically triggered during the early phase of myogenic potentiation, when arteriolar constriction is sensed via physical interactions between arterioles and astrocytic endfeet.

These interactions define a state-dependent system, where vasodynamic and hemodynamic outputs depend not only on current inputs but also on the system’s internal state, shaped by prior activity. In this study, we quantified the first two state-dependent myogenic response modulations and identified the third. Future research can further refine this quantitative framework, and by training it with large multimodal datasets of concurrent brain electrical activity and hemodynamic/vasodynamic signals, lay the foundation for developing a bidirectional predictive model of NGVC. This bidirectional model, which allows for more accurate prediction of vasodynamic or hemodynamic changes based on fluctuations in neural activity and vice versa, has potential applications in early detection of neurovascular diseases [129, 130], implementation of hemodynamic brain-machine interfaces (BMIs) [131], neuroprosthetics [132], functional connectivity studies [133, 134], cognitive neuroscience [135], and pharmacological research [99, 136].

### Novel Methodological Approaches

In this study, we developed both micro-scale (cellular-level) and macro-scale models for arteriolar segments embedded within a coarsely segmented circulatory model of the cerebral vasculature and used that to simulate transient hemodynamic-vasodynamic interactions. Each segment was modeled by transfer functions that relate a wide range of hemodynamic forces—such as WT and WSS—to passive distension and intercellular interactions that adjust the muscular force. The primary goal of introducing these transfer functions was to implement a justifiable autoregulatory mechanism in the developed cerebrovascular model. To add further physiological details, we incorporated appropriate adjustments to the mechanoreactivity of arterioles (by *τ_mct_*, and delays) to represent the range of potential myogenic responses to real-time hemodynamic forces. Ultimately, the interplay between the passive distension and active constriction determined the diameter of the arteriolar segment.

### Vasomotion model

Through computational analysis, we showed that under resting conditions, arteriolar segments cannot maintain a stable balance between passive distension and active constriction, leading to low-frequency, high-amplitude pulsatile dynamics. These spontaneous oscillations serve as key regulatory mechanisms in cerebrovascular physiology, including enhancing tissue perfusion [137, 138], facilitating glymphatic clearance [22, 23], and driving neurovascular synchronization [139]. Therefore, studying the primary signaling mechanisms and the physiological conditions necessary for generating these high-amplitude oscillations is important.

Our proposed vasomotion model differs mechanistically from previously suggested models that rely solely on oscillations generated intrinsically in SMCs via membrane potential or cytosolic calcium oscillators [29, 30]. We showed that while the electrophysiological properties of vascular cells, such as large RMR in aSMC-aEC layers are crucial for the vessel’s excitability to mechanical inputs, the mechanical properties of vascular segments—such as distensibility, mechanosensitivity, and mechanotransduction dynamics—also play indispensable roles in generating oscillations. The positive feedback required to sustain oscillations involves distensibility of vascular segments. Additionally, a highly sensitive mechanosensory subsystem in vessels, capable of translating subtle hemodynamic changes into SMC intracellular signaling, is essential for the proper functioning of the delayed negative feedback in this oscillator. This sensitivity diminishes with increased vascular wall thickness, a common pathological feature in conditions such as hypertension [140]. Furthermore, a mechanotransduction dynamics that is neither overly rapid nor excessively slow yields the highest amplitude of pulsatile dynamics (Fig. 6(b)). In-vivo data suggest that mechanotransduction is slowed in aged mice compared to younger animals [118]. Incorporating these detailed functional and morphological characteristics of cerebral arterioles into the vasomotion model establishes a quantitative framework for investigating key factors contributing to vasomotion attenuation in pathological conditions [30].

### NVC model

In our proposed NVC model, we categorized the vasodilatory effects of neuronal activity on arterioles into two dominant groups: juxtacrine inhibition of the myogenic response by inducing hyperpolarization in SMCs and ECs, and paracrine dampening of the myogenic response via NO production by nNOS-expressing interneurons. We demonstrated that each dampening or inhibition of the myogenic response (excitatory response) is followed by a potentiation (inhibitory response), facilitated by the activity of NPY-expressing interneurons. This computational model could be further refined by elucidating and incorporating the primary mechanisms and relative contributions of other neuronal cell types involved in the CBF regulatory system [141].

Previously proposed NVC models assumed that NPY acts as a direct vasoconstrictor [112, 142, 143]. In the model we proposed, NPY is the facilitator of the myogenic response instead of acting solely as a direct vasoconstrictor. Our rationale was based on the comparisons between our simulation results and reported in-vivo observations. In awake mice, arteriolar vasodynamics in response to instantaneous neuronal activity has a post-stimulus undershoot occurring 6–7 sec after the vasodilation onset, followed by a secondary dilation of smaller amplitude in 10–15 sec post-vasodilation onset [112], similar to the simulation results under *τ*_mct_ = 5 sec as shown in Fig. 10(a). Therefore, it is reasonable to assume that a facilitated myogenic response is the primary driver of both the constriction phase and the secondary dilation. This secondary dilation phase was also observed in our OG stimulation of the somatosensory cortex in ketamine-dexmedetomidine anesthetized mice [8, 114]. In *α*-chloralose anesthetized mice, the constriction phase was smoother, and the post-stimulus undershoot was prolonged, similar to simulated vasodynamics under *τ*_mct_ = 25 sec. It was reported that the pharmacological blockade of NPY-Y1 receptors significantly smoothed the constriction phase [112]. These comparisons suggest that: 1. neuronally evoked SMC NPY1R activation facilitates the myogenic response and, 2. different anesthetic agents have differentiable effects on the dynamics of the myogenic response.

### GVC Model

The astrocytic response in the CBF regulatory system during FH is triggered when astrocyte endfeet sense constriction in arteriolar segments. In-vitro experiments demonstrated that astrocyte endfeet can detect vessel wall strain changes [120], and in-vivo experiments showed that astrocytes contribute to secondary phase of PA vasodilation during sustained FH [12]. Using in-vivo experimental data, our analysis suggests that during sustained FH, astrocytes detect constriction caused by the delayed potentiation of the myogenic response following initial NVC-mediated vasodilation. Astrocytes induce additional vasodilation, which is subsequently followed by another phase of delayed myogenic response potentiation and arteriolar constriction. This constriction is again detected by astrocyte endfeet, triggering another vasodilation. This cycle may occur two to three times during a 30 sec sustained sensory stimulation in awake mice. Notably, in-vivo signals clearly illustrate this stepwise vasodynamic behavior during sustained FH [12].

Despite the frequent astrocytic responses observed in the dynamically functioning mouse brain (Fig. 11(c)), we cannot fully attribute the astrocyte-dependent vasodilatory response to a feedforward, non-metabolic pathway. In 30 sec sustained FH experimental data, astrocyte endfeet Ca^2+^ signaling consistently occurs during the initial constriction phase following the initial NVC-mediated vasodilation, without exception [12].In some published datasets from this same study, subsequent Ca^2+^ signaling events in astrocyte endfeet are not observed. Since neuronal activity did not show a significant decrease over time during their 30 sec FH experiments, it seems likely that the attenuation of astrocytic responses in the later phase of stimulation involved metabolic-related factors. After the first or second step-wise astrocyte-mediated vasodilation in PAs, and the consequent delivery of more blood to the activated region, the astrocyte endfeet may become less preconditioned for substantial Ca^2+^ signaling. These observations suggest that astrocytes likely play a pivotal role in a metabolic feedback mechanism within the brain parenchyma [144]. Elucidating the signaling mediators involved in the astrocytic response in CBF regulation might offer a clearer understanding of their contributions to feedforward and feedback pathways.

We could not find the deterministic feature of astrocytic response in optical or magnetic in-vivo signals acquired from anesthetized mouse brains, including in our experiments, likely because anesthesia disrupts astrocyte calcium signaling [145, 146]. Different anesthetics used in various experimental settings may have different effects on the function of astrocytes [147]. For example, in concurrent recording of BOLD fMRI and Ca^2+^ signaling in anesthetized rat brain during 20 sec sensory stimulations, sometimes post-stimulus glia-related positive BOLD signals were observed immediately post-stimulus, accompanied by increased Ca^2+^ signaling in astrocytes during the stimulus [148]. This post-stimulus positive BOLD fMRI may have arisen from arteriolar vasodilation induced by astrocytic potassium release when vessels initially began to constrict post-stimulus. Astrocytes’ ability to detect vasoconstriction and release potassium into the PVS to induce rapid vasodilation could enhance the amplitude of vasomotion and low frequency vasodynamics in the brain. This mechanism would interrupt the constriction phase early, amplifying vasodilation, which is followed by a more pronounced myogenic response potentiation and vasoconstriction. This idea is supported by in-vivo experimental data which showed that inhibiting astrocyte endfeet Ca^2+^ signaling reduces the amplitude of low-frequency vasodynamics in PAs [120].

### Assumptions and simplifications

In this study, several levels of simplifications and assumptions were incorporated to model an abstract version of the CBF regulatory system, allowing us to focus exclusively on the dominant signaling pathways that shape arteriolar vasodynamics. Despite notable differences between PAs and other vascular segments with large contractility such as sphincters and precapillary arterioles [149–153], it seems plausible that sphincters and precapillary arterioles share similar mechanisms with PAs in shaping their vasodynamics. For example, in the case of vasomotion, sphincters and TZ vessels must also oscillate to provide a similar level of pulsatile dynamics across different cortical depths. If vasomotion were restricted to the arteriolar level and absent in sphincters and precapillary arterioles, capillaries in deeper regions would experience pronounced blood flow oscillations, while blood flow in superficial capillaries would have minimal oscillations. This does not align with physiological expectations. If the NGVC dynamics of sphincters and TZ vessels differ significantly from those of PAs, more blood would be directed toward deeper cortical layers during FH, which again would not align with physiological norms. In-vivo recorded vasodynamics in TZ vessels and Ca^2+^ dynamics in the ensheathing pericytes that encase these vessels corroborate that vasomotion, and low-frequency vasodynamics are also present in TZ vessels, but no vasodynamics have been detected in capillaries during normal brain functioning [154]. It would be valuable to know whether sphincters and precapillary arterioles share similar mechanisms with PAs in regulating CBF.

Previous experimental studies observed delayed vasodilation and increases in positive BOLD fMRI signals in the superficial layers of the awake mouse brain following induced instantaneous neuronal activity [112, 115, 155]. In this study we assumed that there is a cortical depth-dependent time lag in the increase of NO around aSMCs, with superficial aSMCs experiencing this NO increase with a longer delay compared to deeper ones (Fig. 9(d)). The underlying mechanisms behind this phenomenon remain unclear. Given that neuronal activity is known to propagate rapidly across cortical layers, then depth-dependent properties of neurovascular communication become a plausible factor [115]. Further investigation is required to fully elucidate the origins of this delay in superficial layers and to determine whether the CBF regulatory system leverages such depth-dependent kinetics to optimize blood delivery across cortical layers.

It remains unclear whether neurons release potassium around ECs in specific vascular zones and whether ECs in regions with recurrent neural engagement adapt their Kir channel expression in response. This “vascular signaling plasticity” [156] would allow the cerebral vasculature to better match the blood supply with ongoing neuronal activity. In our model, we simplified the EC-dependent mechanism of aSMC membrane potential modulation in response to neuronal activity, considering only an increase in extracellular potassium in aECs during periods of elevated neuronal activity. If a substantial number of ECs across different vascular zones within the parenchyma experience this rise in [*K*^+^]_ex_ during sustained neuronal activity, the endothelial layer could theoretically shift to a quantized state of hyperpolarization and maintain this state throughout the period of elevated activity. This shift would result in more pronounced inhibition of the myogenic response by ECs than modeled in this study when examining vasodynamics in response to step neuronal input. The myogenic response-dependent SMC depolarization following initial NVC-mediated vasodilation could also depolarize aECs through gap junctions and induce a prominent initial overshoot in vasodynamics (Fig. 11(a-right panel)). On the other hand, if local endothelial layer membrane potential is strictly regulated by well-connected EC networks responding to localized neuronal activity, this SMC-mediated EC depolarization may not occur.

### Conclusion and future directions

Our proposed feedforward NGVC model effectively explains most hemodynamic and vasodynamic responses observed in the mouse brain following induced stimuli. However, its limitations become evident with prolonged sensory stimulation in anesthetized mice, where astrocyte involvement is minimal, and NVC mechanisms alone may not fully meet the metabolic demand. In such cases, unexpected biphasic and triphasic CBF increases have been observed during prolonged (64-second) stimulation [77], deviating from the monophasic increase and overshoot expected from a purely feedforward NVC mechanism under anesthesia. We hypothesize that metabolic feedback and adaptive processes, such as RBC deformation in capillaries and K*_AT P_* channel activation in mural cells, may underlie these secondary and tertiary CBF increases, aspects that remain an ongoing focus of our research.

## Methods and Materials

### Hemodynamic-vasodynamic simulations in the cellular-level PA model

To build a computational model for arteriolar segments, the first step is to model how mechanosensitive ion channels in vascular cells are modulated by hemodynamic forces. These forces regulate ion flux through mechanosensitive channels, which, in combination with ion flux through non-mechanosensitive channels, shape the electrochemical gradients and resting conditions of the cell. To define the segment-specific dynamic range of hemodynamic forces, we extracted *WT*_max_ and *WSS*_max_ for each arteriolar segment. This range was determined under the assumption of static autoregulation in the cerebral vasculature, where each segment operates under specific WT-MT and WSS-MT transfer functions, enabling autonomous adjustment of vessel segment contraction state based on hemodynamic inputs. Given that WT-MT has more control over MT adjustments than WSS-MT within the physiological range of hemodynamic forces [34], we particularly aimed to build a more detailed model of aSMCs than aECs in this study.

The SMC electrophysiology was modeled using a system of ODEs, primarily adapted from Karlin’s study [72], with adjustments for PA SMCs. The T-type VOCC/BK/RyRs microdomain, which minimally affects PA SMC negative feedback in MT regulation, was not incorporated in the model, while a BK-Glu microdomain was included as part of the NGVC mechanisms. This microdomain incorporates NMDA receptors that, upon activation by neuron-released glutamate, facilitate Ca^2+^ efflux to activate BK channels. The inclusion of such microdomains reflects the requirement for localized [Ca^2+^] to exceed the cytoplasmic [Ca^2+^] in SMCs to activate Ca^2+^-sensitive channels. Additionally, a wall tension-transducing microdomain (WT-TM) was incorporated to account for the Ca^2+^-dependent activation of key channels involved in MT regulation, such as TMEM16A and TRPM4. Without the WT-TM, a positive feedback loop would occur, where depolarization triggers further Ca^2+^ influx via VOCCs, continuously activating these Ca^2+^-sensitive depolarizing channels. Thus, incorporating a macrodomain where Ca^2+^ modulation is controlled exclusively by non-voltage-gated channels, such as mechanosensitive TRPC6, was essential.

For the mechanosensory subsystem of the SMC model, given that the signaling cascades associated with this subsystem are not fully understood, we applied simplifications that may not be entirely physiologically accurate. We assumed that WT linearly influences phosphatidylinositol 4,5-bisphosphate (PIP2) concentration through the mechanoactivation of G-protein-coupled receptors (GPCRs) [157]. Specifically, an increase in WT enhances the conversion of PIP2 into IP3 and DAG. This relationship was modeled as a quasi-linear sigmoidal relationship between increased WT and reduced PIP2, leading to both a decrease in the probability of Kir channel activation [157] (Fig. 5(b)) and, in parallel, an increase in DAG and IP3 concentrations (Fig. 3(a)). The elevated DAG and IP3 subsequently raise the open probability of TRPC6 channels [39] and IP3 receptors (IP3Rs) within the WT-TM, respectively. The activation of TRPM4 channels in the WT-TM, via transient Ca^2+^ release from IP3Rs, generates transient inward cation currents (TICCs), where the frequency of TICCs is influenced by the concentrations of Ca^2+^ and IP3 within the WT-TM. In our model, this cascade was simplified by focusing on steady-state conditions. This steady-state approach streamlines the system of ODEs by eliminating the need to model numerous microdomains where transient currents occur. Instead, the cumulative effects of these transient currents are captured in the steady-state activation levels of channel pools, which reduce the computational demands for simulations. With this simplification, we assumed that IP3 concentration directly increases the open probability of TRPM4 channels rather than acting indirectly through IP3Rs [68]. As a result, the activity of TRPM4 channels is regulated by WT/[DAG]/TRPC6/[Ca^2+^]_TM_ and WT/[IP3]_TM_. Similarly, the mean ion flux through TMEM16A channels, calcium-activated chloride channels assumed to be localized in the WT-TM, is regulated by WT/[DAG]/TRPC6-[Ca^2+^]_TM_ and the SMC membrane potential. The electrophysiology of the developed model was subsequently validated by simulating the behavior of the SMC under various conditions, including voltage-clamp experiments (Fig. 3(d)) and pressure myography maneuvers (Fig. 3(b)). For detailed equations governing SMC electrophysiology, including those describing channel currents and intracellular ion concentrations, readers are referred to [72], and to our code implementation.

Upon establishing and validating the SMC electrophysiological model, we integrated it with another critical component of the arteriolar segment model: vessel contraction adjustments by modeling the interplay between passive distension and SMC constriction mechanisms. Importantly, its output (i.e. vessel diameter) serves as the primary input for hemodynamic analysis. To effectively represent the SMC constriction mechanism through the dynamic modulation of MLCK and MLCP activities, we focused on the dominant signaling mechanisms proposed in this study. Specifically, the model links the SMC electrophysiology framework via Ca^2+^-mediated modulation of MLCK, while enabling NO/cGMP/PKG-driven modulation of MLCP activities in response to external stimuli from ECs and nNOS-expressing interneurons. To achieve this, we employed the four-state Michaelis-Menten kinetics model proposed by [85] and adapted by [87]. This model captures transitions between four distinct myosin states: unphosphorylated myosin (M), phosphorylated myosin (Mp), actin-bound unphosphorylated myosin (AM), and actin-bound phosphorylated myosin (AMp). A system of ODEs describes the temporal evolution of these states, with rate constants defining the transitions. The Ca^2+^ dependency of actomyosin interactions, along with the effects of NO/cGMP/PKG signaling, was incorporated by modulating these rate constants. By conserving the total myosin across all states, the model provides a robust framework for simulating SMC constriction dynamics. For further details on the implementation and parameterization of this model, readers are referred to [158], and to our code implementation.

The PA segmental model was then integrated into the cerebrovascular model. Algorithm 1 outlines the hemodynamic-vasodynamic simulation workflow developed to investigate the kinetics of vasomotion and FH dynamics using this cellular PA model. The hemodynamic analysis was based on the research conducted by Secomb’s [159] and Pries’s [35] groups.

### Hemodynamic-vasodynamic simulations in the network of macro-scale models of coupled arteriolar segments

The developed simplified macro-scale model of arteriolar segments incorporates the delayed myogenic response, passive distension, and electrical coupling between adjacent segments. Algorithm 2 outlines the simulation workflow. The model begins by setting initial conditions, calculating *WT_avg_* based on IP, D, and constant vessel wall thickness (h). The variables related to the myogenic response, passive distension, and electrical coupling in the model are defined to simulate vasomotion and FH dynamics. In the analysis of vasomotion dynamics, where the myogenic response is not modulated by stochastic *γ*-band signals, the membrane potential of each arteriolar segment is initially computed solely based on its hemodynamic inputs (*V_init_*(*AS*)). To model the resistive-capacitance nature of electrical coupling between segments, a secondary coupling loop is introduced. This loop operates at a finer time step (*dt*_2_ = 0.4 msec) to avoid fast and large influences of adjacent segments on each other. In this loop, the difference in membrane potential between adjacent segments is calculated, and the membrane potential of each segment is iteratively updated. Δ*V*_coupling_(*AS, j*) represents the cumulative voltage differences between each segment and its adjacent segments. The final membrane potential at the end of this loop (*V_final_*(*AS, i*)) is used for subsequent calculations in the main loop. The passive portion of the diameter (*D*_PD_(*AS, i*)) is then computed based on the real-time WT and depends on the distensibility parameters and the minimum diameter under no pressure conditions (unpressurized state), which was assumed to be 90% of the maximum active diameter (Fig. 3(b)). The total diameter of each segment is the sum of the MR portion and the passive portion. Finally, if the updated diameter exceeds the maximum allowable diameter change in one main loop iteration, the change is constrained by the maximum allowable dilation (Δ*D*_maxdila._ (*AS*)) or constriction (Δ*D*_maxconst._ (*AS*)) rates. During the FH vasodynamic analysis, all steps remain unchanged except for the inhibition of MR by the *γ*-band signal (*GB*), and the dampening of MR by the smoothed *γ*-band (*GB_S_*), as described in Section 3.

#### Algorithm 1 hemodynamic-vasodynamic simulation workflow in cellular-level PA model

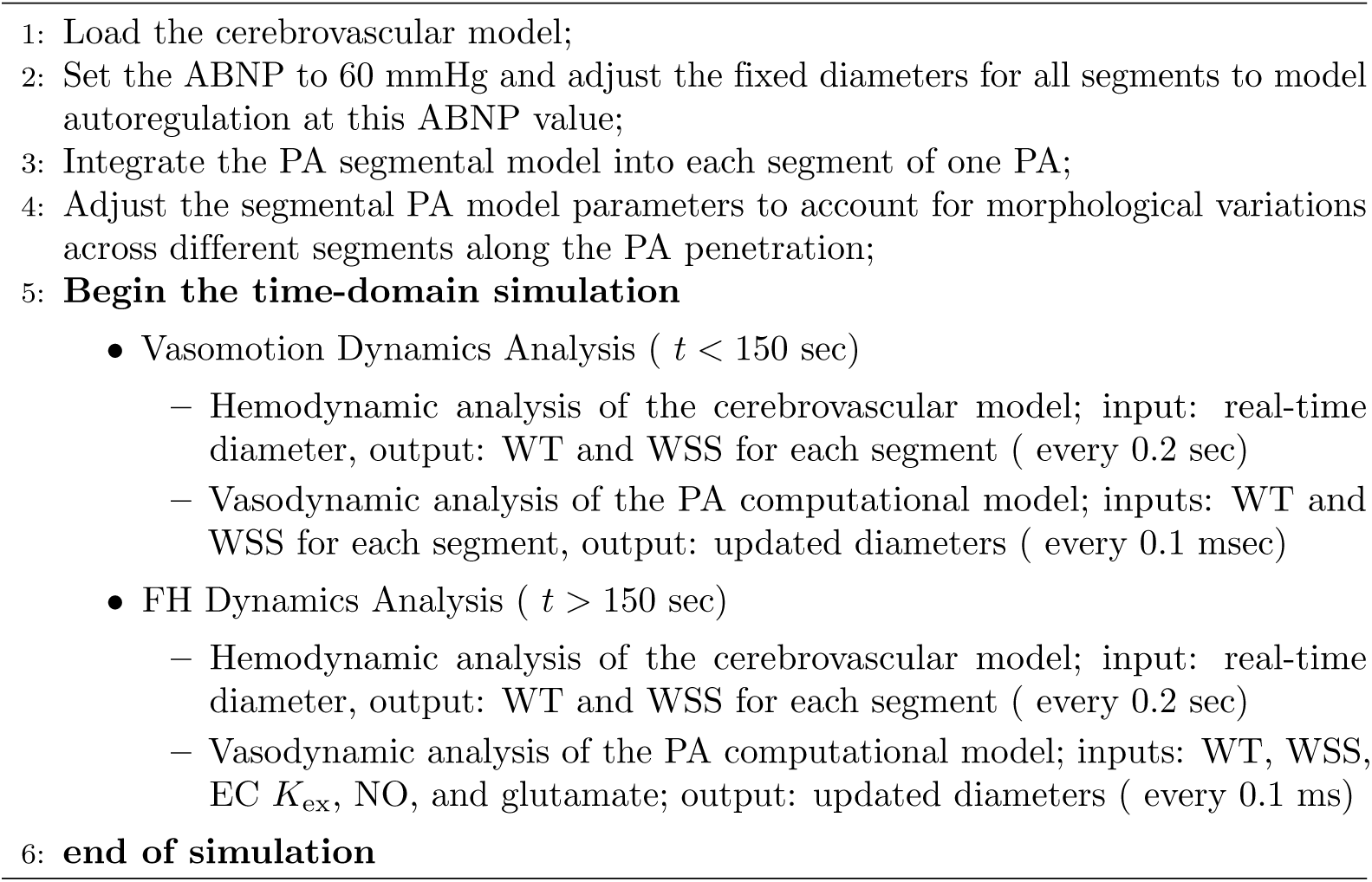

### Animal Preparation and In-Vivo Imaging

All animal procedures were approved by the University of Wisconsin Institutional Animal Care and Use Committee (IACUC). We used 6–12 month-old transgenic optogenetic mice (THY:ChR2/H134R-YFP). Mice were anesthetized with isoflurane (1.5%–2.0% in oxygen), and the scalp was retracted to thin the skull over the somatosensory cortex. A 3 mm coverslip was installed approximately 1.5–2 mm lateral and 1.5–2 mm posterior to bregma. The average heart rate during surgery was 450–550 bpm. Afterward, anesthesia was switched to ketamine (25–50 mg kg^-1^ SC) and dexmedetomidine (0.05–0.5 mg kg^-1^ SC) for imaging. Heart rate during imaging was 250–350 bpm, with peripheral capillary oxygen saturation between 95–100%. Body temperature was maintained at 36.5–37.5°C using a heating pad. Cerebral blood flow (CBF) was recorded using laser speckle contrast imaging (LSCI; RFLSI III, RWD Life Sciences).

### Statistical Analysis

In Section 3, we quantified the onset time of OG-evoked vasodilations at various cortical depths from Uhlirova’s dataset to assess the speed of vasodilation propagation in response to instantaneous neuronal activity. A sigmoidal function was fitted to the diameter increase data to determine the onset of dilation, identified by the point where the first derivative of the fitted curve crossed a predefined threshold, marking the start of the diameter increase.

#### Algorithm 2 hemodynamic-vasodynamic simulation workflow in a network of macro-scale models of coupled arteriolar segments

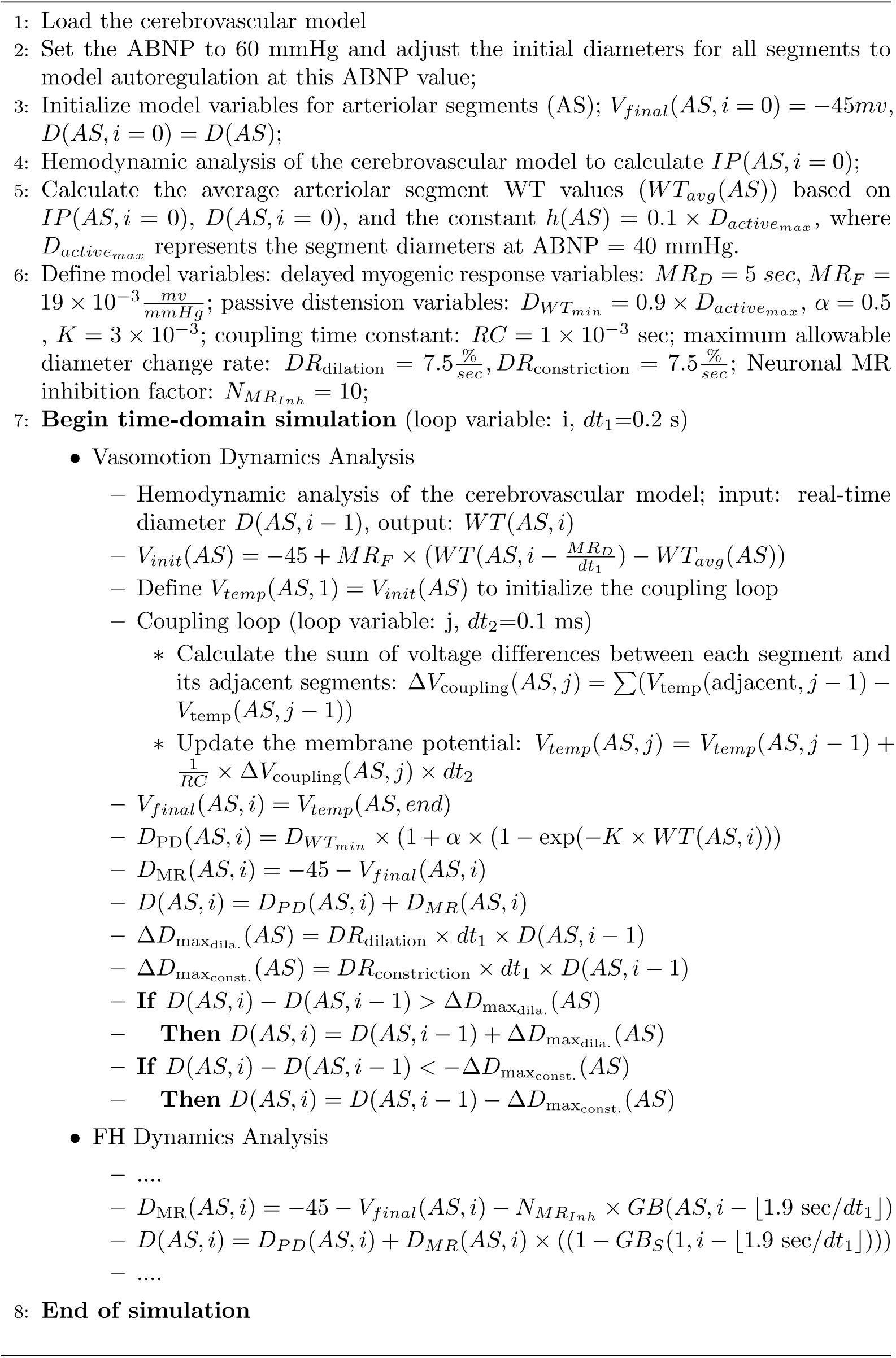

## Acknowledgments

This study was conducted using resources and facilities at the William S. Middleton Memorial Veterans Hospital,

